# Epigenetic conditioning improves sequence-based modeling of gene regulation across cell types and alleles

**DOI:** 10.64898/2026.06.02.729723

**Authors:** Oberon Dixon-Luinenburg, Ayesha Bajwa, Mitchell R. Vollger, Andrew B. Stergachis, Aaron Streets, Nilah M. Ioannidis

**Affiliations:** UC Berkeley–UCSF Graduate Program in Bioengineering, University of California, Berkeley, Berkeley, CA, USA; Center for Computational Biology, University of California, Berkeley, Berkeley, CA, USA; Department of Electrical Engineering and Computer Sciences, University of California, Berkeley, Berkeley, CA, USA; Department of Human Genetics and Utah Center for Genetic Discovery, University of Utah, Salt Lake City, UT, USA; Division of Medical Genetics, Department of Medicine, University of Washington, Seattle, WA, USA; Department of Genome Sciences, University of Washington, Seattle, WA, USA; Brotman Baty Institute for Precision Medicine, Seattle, WA, USA; Department of Bioengineering, University of California, Berkeley, Berkeley, CA, USA; Biohub, San Francisco, CA, USA; Department of Applied Mathematics, University of California, Santa Cruz, Santa Cruz, CA, USA

**Keywords:** genomic deep learning, epigenetics, CpG methylation, conditioning, sequence-to-activity, long-read sequencing

## Abstract

Epigenetic state modulates gene regulation in a manner not always predictable from DNA sequence alone, yet current genomic deep learning models do not leverage epigenetic state as input. We present MethylSeqNet, a model that conditions pretrained sequence embeddings on CpG methylation, a stable epigenetic mark increasingly available from long-read sequencing data. Using a novel conditioning mechanism enabling scalability and interpretability, MethylSeqNet improves predictions in cases where differential epigenetic state drives regulatory variation. We show improvements over a sequence-only baseline for cell-type-specific chromatin accessibility and transcription. Epigenetic conditioning enables prediction of phenomena not encoded in allele sequence, including parent-of-origin imprinting, random monoallelic activity, and X-inactivation. We highlight a promising application of methylation conditioning by predicting the effects of a structural rearrangement in one rare disease patient case study. In silico motif insertion analysis confirms that MethylSeqNet learns methylation-dependent regulatory grammar, establishing a paradigm for integrating epigenetic information into genomic deep learning with immediate applications in rare disease interpretation.

## 1 Introduction

Accurate prediction of regulatory activity from DNA sequence across diverse human cell types, cell states, and haplotypes is of broad importance for diagnosis, treatment, and understanding molecular mechanisms of disease. Furthermore, generalizable predictive models can be used to better understand the sequence grammar of the regulatory genome. In the past decade, supervised deep neural networks trained on large corpora of molecular phenotype measurements have shown success in predicting sequence-driven *cis*-regulatory activity across a range of cell types and shown promise in predicting disease-relevant variant effects [1–8].

However, state-of-the-art sequence-to-activity models do not currently consider information beyond primary DNA sequence when making predictions. Long-read sequencing technologies, which are increasingly being applied to profile human genetic diversity and study undiagnosed genetic diseases [9, 10], natively measure DNA modifications along with primary sequence. CpG methylation is the most common and biologically relevant DNA modification in vertebrates and forms a key part of persistent epigenetic state. Epigenetic state can directly drive differential regulatory activity such as in X-inactivation, imprinting, and epigenetic silencing [11–13]. It can also be a strong signal for sequence-driven regulation, as with deterministic epigenetic programming differences between cell types [14] and sequence variant regulatory effects [15]. Prior models establish that CpG methylation exhibits sequence-context-dependent dynamics [16–18] and can in some cases be predicted from sequence variants [19]. However, current models do not leverage the increasing availability of CpG methylation data to condition sequence-to-activity predictions on epigenetic state.

We present MethylSeqNet, a deep neural network built using a novel sequence-to-activity framework that conditions predictions on epigenetic state. Because training data that couples CpG methylation with functional activity remains limited, MethylSeqNet is designed to leverage a pretrained sequence encoder. Conditioning is applied through a novel mechanism that we call factorized representation modulation (FaRM). FaRM enables interpretation of sequence element methylation sensitivity while achieving efficient training and inference when conditioning a high-dimensional pretrained sequence embedding on a scalar epigenetic state. MethylSeqNet is trained on both cell-type-differential and haplotype-differential data.

Across all prediction tasks, MethylSeqNet achieves competitive or improved prediction performance when compared to a parameter-matched unconditioned sequence baseline. We observe large improvements over an unconditioned sequence baseline where differential epigenetic state drives differential activity at the same sequence elements. We also see improvements in cases where differential activity may be driven by sequence variants, but altered epigenetic state provides a stronger predictive signal to the model. As biologically expected, we see the largest improvements in cases of allelic imbalance that are not deterministically encoded in the sequence of the alleles themselves, such as X-inactivation. We demonstrate that MethylSeqNet can generalize to correctly predict disease-associated allelic imbalance in an undiagnosed disease patient with a balanced translocation. Finally, using in silico motif insertion experiments, we computationally validate that MethylSeqNet has learned a rich grammar of transcription factor (TF) motifs with methylation-conditional activity, and that this learned sensitivity aligns with orthogonal characterization of these TFs. By demonstrating how epigenetic state can be leveraged to substantially improve sequence-based predictions of regulatory activity, MethylSeqNet establishes a general framework for integrating epigenetic information into genomic deep learning models, with immediate applications in interpreting patient-specific regulatory variation from long-read sequencing data.

## 2 Results

### 2.1 MethylSeqNet improves regulatory activity prediction by conditioning on methylation

The MethylSeqNet architecture (Fig. 1a) processes input DNA sequence and CpG methylation through sequential modules described in Eq. 1 to predict activity **ŷ**. First, DNA sequence **x** passes through a pretrained sequence encoder *ϕ* (Eq. 1a) to produce sequence embedding **h**. Next, **h** and the conditioning state **s**, representing local CpG methylation level across genomic bins, pass through an epigenetic conditioning mechanism *g* to produce conditioned representation **z**, encompassing both sequence and methylation (Eq. 1b). Finally, an output head *ψ* takes in **z** to predict chromatin accessibility and gene expression levels in many cellular contexts (Eq. 1c).

**Fig. 1.**
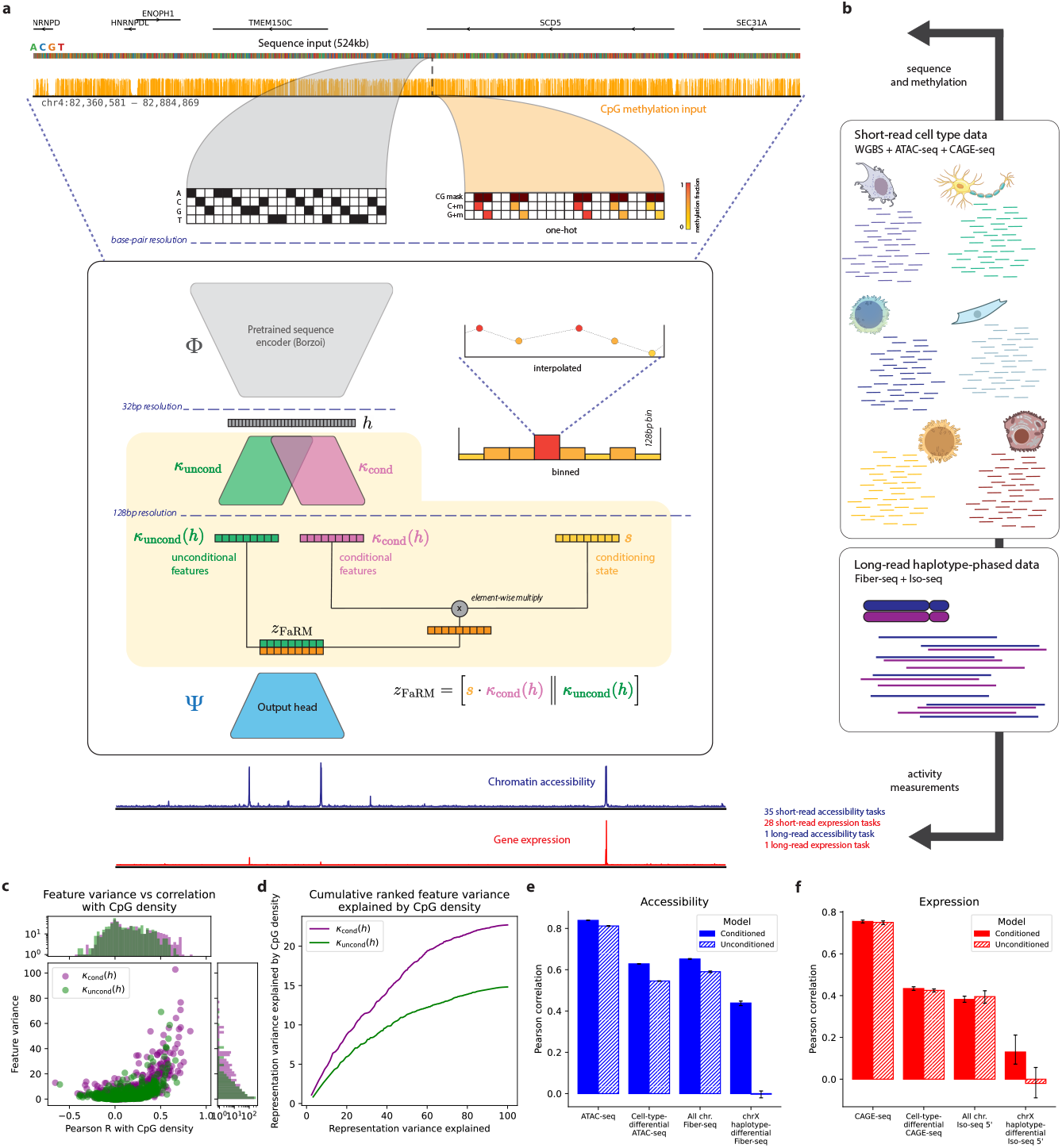
MethylSeqNet predicts chromatin accessibility and gene expression from sequence and methylation. **a**, The MethylSeqNet architecture consists of a pretrained encoder, a methylation-conditioning mechanism, and a learned output head with a final layer containing a filter for each task. MethylSeqNet maps from sequence and methylation input to accessibility and expression predictions. **b**, MethylSeqNet’s training data consists of mostly short-read data across many distinct cell types (35 accessibility tasks, 28 expression tasks) as well as long-read haplotype-phased data (1 accessibility task and 1 expression task). **c**, Scatterplot of feature variance and neural network activation correlation with CpG density for conditional and unconditional features. **d**, The cumulative ranked feature variance explained by CpG density for conditional and unconditional features. **e**,**f**, Comparison of conditioned vs unconditioned MethylSeqNet predictive performance on the test set for subsets of accessibility and expression tasks focused on cell-type differences (ATAC-seq and CAGE-seq). 95% confidence intervals from 100 bootstrap resamples (N = full test set size, with replacement) are shown.

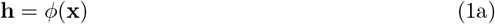

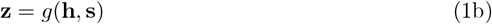

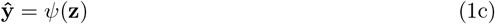

This architecture was inspired by factorized regulatory sequence models [4, 20], which provide interpretability via architectures designed to separate information into different pieces of the learned representation, and by successful applications of transfer learning to large pretrained models of DNA regulation [21, 22], which can achieve strong performance even in low-data regimes.

We propose a novel conditioning mechanism designed to apply epigenetic conditioning while maintaining the interpretability of the learned representation. Factorized representation modulation (FaRM) maps **h** and **s** to **z** by learning a partitioning of **h** and applying the conditioning signal directly to one partition through element-wise multiplication (Eq. 2). We define

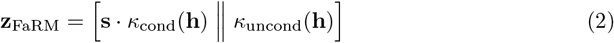

where *κ*_cond_ and *κ*_uncond_ are learned nonlinear transformations of **h** and [· ‖ ·] denotes concatenation. Assuming the training distribution covers sufficient conditioning state diversity for a given sequence, FaRM separates methylation-dependent and methylation-independent sequence features by construction.

We trained MethylSeqNet jointly on 63 short-read cell type tasks and two long-read haplotype-phased tasks (Fig. 1b). We matched cell types from whole-genome bisulfite sequencing (WGBS) [14], pseudobulked single-cell assay for transposase-accessible chromatin (scATAC-seq) [23], and cap analysis of gene expression (CAGE-seq) [24] primary cell atlases according to Supplementary Table 1. Thus, our training data captures a range of *cis*-regulatory patterns and epigenetic states across different *trans*-regulatory environments. The haplotype-phased data, sequenced using Fiber-seq and PacBio Iso-seq [10, 25, 26], allowed us to directly train on allelic imbalance, capturing differential activity at the same loci with identical *trans* environment driven partly or entirely by epigenetic differences. By training on distributions across cell types and distributions across haplotypes, we aimed to capture their joint properties in our model, allowing us to probe phenomena such as allelic imbalance in cell types for which the model’s training data did not contain allelic imbalance. Following standard practice in supervised sequence modeling, we learned these tasks in a multitask framework, implicitly learning the *trans* environment through task-specific parameters in the final layer.

On the joint cell types and haplotypes data, we first demonstrated that augmenting sequence input with CpG methylation in a small convolutional neural network (CNN) jointly processing **x** and **s** drives clear improvement in the predictive performance (Supplementary Fig. 1a, Methods). Smoothing the methylation landscape does not degrade the performance of a model until done at scales in excess of 1 kb (Supplementary Fig. 1b), as we expect from the biological correlation of CpG methylation state at nearby dinucleotides [27]. We therefore handle CpG methylation conditioning in 128 bp bins, a typical resolution for molecular activity predictions, throughout the rest of this study.

We achieved strong prediction performance and rapid convergence by applying transfer learning utilizing Borzoi [6] as our pretrained sequence encoder *ϕ* in Eq. 1a. We applied the binned conditioning state with a five-layer CNN output head to integrate methylation from an 8 kb receptive field for each prediction bin. FaRM shows similar performance to other conditioning mechanisms (Supplementary Fig. 2a, Methods) but with two primary advantages. First, FaRM conditional representations can be interpreted as linear sensitivity to the scalar conditioning variable, facilitating biological interpretation. Second, FaRM is computationally efficient in the many-conditioned-state case, requiring 27% less memory and running training steps 40% faster for the same number of features compared to FiLM (Supplementary Fig. 2b). We gain additional speed and memory improvements because, by separating Eq. 1a and Eq. 1b, the sequence encoder requires only one forward pass per sequence and shares the embedding across many conditioning states, rather than jointly processing **x** and **s** with all model parameters.

While MethylSeqNet primarily predicts accessibility and expression outputs, our framework also supports imputation of the conditioning methylation state as another prediction output. The resulting imputed methylation shows strong concordance with measured methylation, including for cell-type-differential methylation (Supplementary Fig. 2c). Using imputed methylation from a MethylSeqNet model trained with conditioning methylation only slightly reduces performance compared to a model trained without conditioning (Supplementary Fig. 2d). Furthermore, imputed methylation can help interpret failure modes for unconditioned sequence models, identifying cases where methylation conditioning provides the most advantage. For conditioned vs unconditioned activity predictions throughout this study, we use MethylSeqNet models trained with and without conditioning respectively.

To establish that factorized representations diverge from one another, we explored neural network activation correlation with CpG density, since the presence of CpGs within a TF binding motif suggests methylation-sensitivity [28]. While both conditional and unconditional representations contain features with high correlation to CpG density, the conditional representation is enriched for higher correlations (Fig. 1c). CpG-correlated features have higher variance across the test set than non-CpG-correlated features, meaning they can contribute more to differential predictions across loci. We computed the total representation variance explained by CpG density for both conditional and unconditional representations and found that the variance explained was a factor of two larger for the conditional representation (Fig. 1d). We found the higher CpG density correlation in the conditional representation is substantially strengthened by the inclusion of allelic imbalance data in training, indicating better partitioning of the methylation-sensitive features (Supplementary Fig. 2e). However, CpG density explains only 22% of the variance, meaning the full encoding of the input DNA sequence contains more information than CpG density alone. Results 2.5 further explores the sequence elements driving conditional neural network activations.

Compared to a parameter-matched sequence-only output head, we see clear improvements in prediction performance across the test set for both cell type (Fig. 1e) and allelic imbalance (Fig. 1f) data. Improvements are larger for chromatin accessibility than for gene expression and largest for differential predictions between cell types or haplotypes rather than for peak predictions within a cell type or haplotype. For cell-type-differential activity, a learned mapping from methylation to activity underperforms unconditioned predictions, and both underperform a conditioned MethylSeqNet model (Supplementary Fig. 2f). Therefore, the performance boost from conditioning relies on the combination of sequence and methylation information and cannot be recapitulated from either alone.

### 2.2 MethylSeqNet improves cell-type-specific activity prediction

We examined MethylSeqNet’s test set performance across individual cell types and loci to characterize improved prediction performance. ATAC-seq prediction for almost every cell type showed an improvement in Pearson correlation between predictions and measurements across the test set compared to unconditioned predictions (Fig. 2a). While CAGE-seq improvements were minimal for peak identification (Supplementary Fig. 3a), the improvement was substantially larger for cell-type-differential activity (Fig. 2b, Supplementary Fig. 3b, Methods). This result is consistent with our expectation that methylation conditioning is more important for sequence elements with varying epigenetic states as opposed to those whose epigenetic states are consistent across the training distribution; the additional signal provided by methylation is less informative when the methylation state of a sequence element is the same for all training samples.

**Fig. 2.**
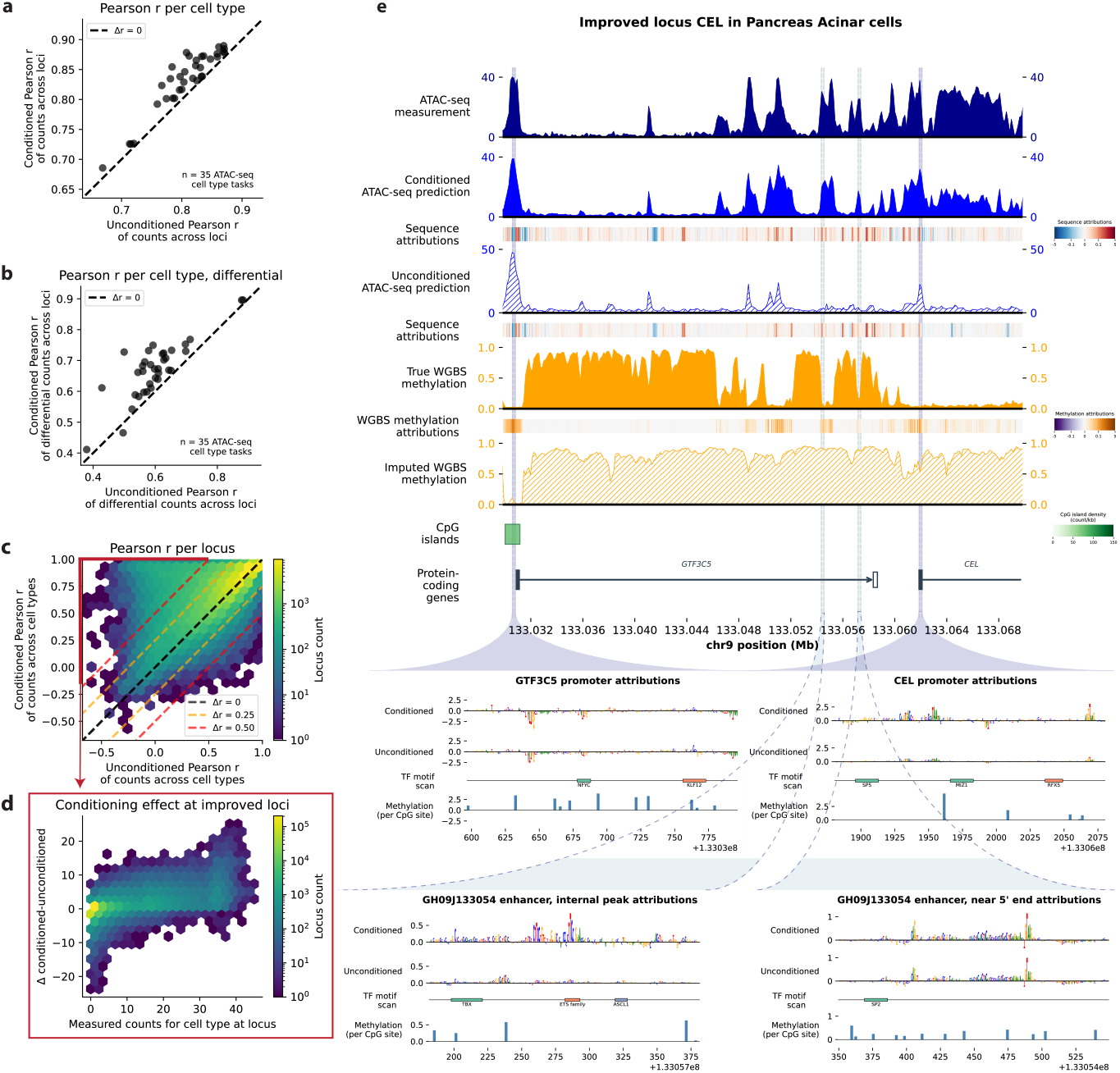
MethylSeqNet improves cell-type-specific activity predictions. **a**, Scatterplot comparison of MethylSeqNet conditioned vs unconditioned prediction-measurement Pearson correlation for ATAC-seq test set by cell type. Each point represents one cell type/task, and the dashed line indicates equal performance. **b**, As in **a**, but for cell-type-differential Pearson correlation calculated from the difference from mean activity across cell types for the input sequence (Methods). **c**, Hexbin plot comparison of conditioned and unconditioned test set prediction performance across cell types for all loci. Each point represents a locus, and the central dashed line indicates equal performance. **d**, Hexbin plot with one count for cell type from each locus in **c** with *>*0.5 Pearson improvement plotting differential conditioned minus unconditioned predictions against measured counts. **e**, Visualization of genomic tracks at the *CEL* locus in pancreatic acinar cells. Tracks display (from top to bottom) accessibility measurements, conditioned and unconditioned ATAC-seq predictions from MethylSeqNet along with their respective sequence attributions calculated using integrated gradients, WGBS methylation measurements, MethylSeqNet imputed WGBS methylation, methylation attributions calculated using integrated gradients, and CpG island and gene annotations. Magnified inserts show sequence and methylation attributions for two promoters (dark blue) and two regions of a large enhancer (light blue).

We investigated three hypotheses for why some cell types underperform others. First, different cell types have varying WGBS read depth, resulting in some noisier conditioning states. Second, cell types from the different atlas datasets vary in their CAGE-seq and ATAC-seq match quality to the WGBS cell types. Finally, the Borzoi model did not see all the MethylSeqNet cell types in pretraining. We show that WGBS read depth and cell type matching explain some, but not all, of the differences in ATAC-seq (Supplementary Fig. 3c) and CAGE-seq (Supplementary Fig. 3d) performance. However, comparing Borzoi as a pretrained sequence encoder to a less powerful sequence encoder trained from scratch, we observe tightly correlated prediction improvements by cell type for ATAC-seq (Supplementary Fig. 3e). CAGE-seq shows a weaker relationship (Supplementary Fig. 3f), but for the CAGE-seq tasks these are all within the Borzoi training corpus. We conclude that the performance difference not explained by WGBS read depth and cell type matching is likely intrinsic to the cell types or the individual ATAC-seq and CAGE-seq measurements, rather than the pretraining distribution. We hypothesize that some cell types may contain a larger set of peaks with unique sequence grammar that is hard to predict without either conditioning or more training data.

Because CAGE-seq tasks show less improvement from methylation conditioning, the rest of this section focuses on ATAC-seq improvements. We examined all test set loci with accessibility peaks in at least one cell type. The Pearson correlation of predicted vs measured activity across cell types at each locus in the conditioned vs the unconditioned model (Fig. 2c) shows that test-site-wide improvements emerge from a subset of loci: Pearson correlation across cell types improves by over 0.25 for 26,937 peaks, worsens by over 0.25 for 1,850 peaks, and remains within a ± 0.25 range for the remaining 126,801 peaks. 9,861 of the 26,937 improved peaks see a Pearson correlation improvement of over 0.5 (Supplementary Fig. 3g).

For these most improved loci, we examined the differential conditioned vs unconditioned predictions by locus in each cell type. From this, we identified two distinct dynamics (Fig. 2d). First, there are loci for which the activity in one or more cell types is overpredicted by the unconditioned model and conditioning helps recognize silencing. Second, there are loci for which the activity in one or more cell types is underpredicted by the unconditioned model and conditioning helps recognize activation.

We show a representative example of improved prediction around the well-characterized *CEL* gene (Fig. 2e), which we expect to be active in only our pancreatic acinar cell type [29]. Examining the trace, we see strong accessible peaks upstream and downstream from the transcription start site (TSS), including several within the enhancer GH09J133054 which is associated with *CEL* and belongs to the super-enhancer SE 47567, active in pancreas [30]. The true methylation landscape contains a large hypomethylated region extending across the gene body and several sites upstream. In contrast, the unconditioned sequence model predicts a methylation landscape with mostly high methylation levels and some localized regions of lower methylation. The large-scale hypomethylation may be driven by a few strong TF binding events at specific loci during differentiation or by relatively distal elements within the SE 47567 super-enhancer which then recruit machinery to program the larger region. This sort of regulatory dynamic, wherein local sequence signals are weak, could explain why the unconditioned sequence model systematically underpredicts accessibility at many loci within 15 kb of the *CEL* TSS. We visualize input attributions (Fig. 2e) to establish the relative contributions of specific loci to the predictions, with separately calculated sequence and methylation attributions enabling fine-grained interpretation (Methods). Sequence attributions reveal that conditioning allows MethylSeqNet to attend more strongly to sequence elements near the TSS and within GH09J133054. Methylation attributions illustrate that different parts of the epigenetic landscape vary in importance for driving activity. We further examine the learned importance of specific sequence elements and their methylation state in Results 2.5.

We hypothesized that highly improved loci may reflect propagating epigenetic state or distal interactions, such as within chromatin compartments [31]. However, relative enrichment in A compartments (active) vs B compartments (silenced) did not show significant differences between active and silenced subsets of improved-by-conditioning loci (Supplementary Fig. 3h). We investigated four additional examples of highly improved loci and identified that the unconditioned model often fails to accurately predict local methylation at the incorrectly predicted peaks, whether overpredicted or underpredicted (Supplementary Fig. 4a-d). Thus, while anecdotal evidence suggests MethylSeqNet conditioning improvements may derive from epigenetic state reflecting long-range interactions, it is unclear whether this is the dominant source of improved performance. Other possible explanations include the missed sequence elements simply being, in and of themselves, more distinct from examples in the training set. For example, some activity is driven by multi-TF interactions that are insufficiently abundant across the genome to be learned by current supervised models without additional data, such as activity measurements for which TF interactions are actively perturbed.

### 2.3 MethylSeqNet learns allelic imbalance from training on haplotype-phased data

Allelic imbalance is typically harder to predict from sequence alone than is differential activity across healthy cell types, which derives largely from cell-type specific sequence motifs. While allelic imbalance that is driven by sequence variants may be predictable from sequence, much of the strongest allelic imbalance present in the genome is driven by epigenetic effects that differ across haplotypes: X-inactivation, parent-of-origin imprinting, and random silencing impact thousands of genes and tens of thousands of elements [32–34].

We inspected whether conditioned MethylSeqNet improves functional activity prediction on the test set haplotype-phased allelic imbalance data compared to unconditioned sequence modeling, which sees sequence differences but not the distinct epigenetic states of the two haplotypes. Our results show that haplotype-phased chromatin accessibility (Fiber-seq) and gene expression (Iso-seq) are generally better predicted when conditioning on methylation, compared to the unconditioned baseline (Fig. 3a,b). As seen for the equivalent short-read unphased measurements, the improvements are substantially greater for accessibility than for expression. For haplotype-differential activity across all loci, Fiber-seq prediction performance is relatively strong with conditioning, whereas without conditioning the model cannot do better than random chance. While unconditioned sequence models can predict eQTL effects better than chance, here we include all loci in the test set and all phased SNP variants across chromosome X and the autosomes respectively. For Iso-seq predictions, only chromosome X is better than chance with methylation conditioning: the relative noise levels for the phased Iso-seq are likely swamping the signal from the relatively few strongly differentially expressed autosomal genes. With higher-read-depth data from more cell lines or primary cells, we posit that the MethylSeqNet framework should be able to achieve performance boosts at least analogous to those seen for CAGE-seq.

**Fig. 3.**
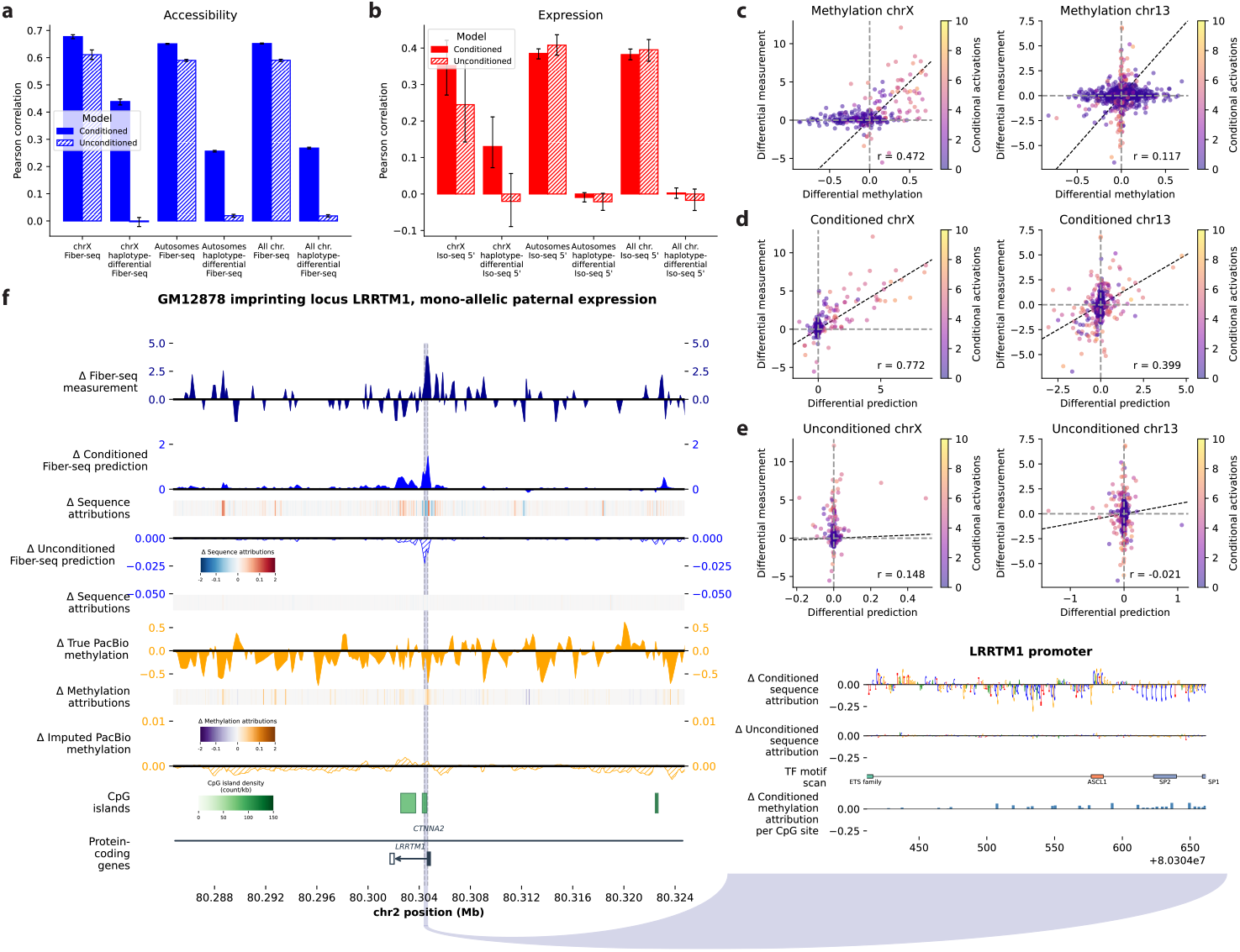
MethylSeqNet conditioning improves allelic imbalance prediction. **a**,**b**, Comparison of conditioned vs unconditioned MethylSeqNet predictive performance on the test set for subsets of accessibility and expression tasks focused on capturing allelic imbalance (Fiber-seq and Iso-seq). 95% confidence intervals from 100 bootstrap resamples (N = full test set subset size, with replacement) are shown. **c**, Scatterplots of differential accessibility measurements against differential methylation between haplotypes for chromosomes X and 13. Points are colored by neural network activations of methylation-conditional features and dashed lines indicate y=x (black) and x=0, y=0 (gray). **d, e**, Scatterplots of differential accessibility measurements against differential accessibility predictions between haplotypes for chromosomes X and 13 in the conditioned and unconditioned models. **f**, Example of one imprinting locus where MethylSeqNet improves accessibility prediction: *LRRTM1* in GM12878 lymphoblastoid cells. We visualize haplotype-differential genomic tracks for accessibility measurements, conditioned and unconditioned predictions from MethylSeqNet along with their respective sequence attributions calculated using integrated gradients, PacBio methylation measurements, MethylSeqNet imputed PacBio methylation, methylation attributions calculated using integrated gradients, and CpG island and gene annotations. We zoom in to show sequence and methylation attributions for the *LRRTM1* promoter.

To demonstrate MethylSeqNet on allelic imbalance in TSS activity across genes, we first show that haplotype-differential methylation alone is insufficient as a predictor of differential activity for chromosomes X and 13 (Fig. 3c). While differential activity is typically either driven by or correlated with differential epigenetic state, sequence elements have varying sensitivity to methylation that is missed without knowledge of specific sequence context. Next, we plot MethylSeqNet haplotype-differential chromatin accessibility predictions and measurements across chromosomes X and 13 and observe a roughly linear relationship (Fig. 3d). Chromosome X has much stronger allelic imbalance but MethylSeqNet predictions correlate well on both. Unconditioned sequence predictions, however, capture essentially none of the differential activity (Fig. 3e). On chromosome X, the copy silenced by X-inactivation cannot be predicted from sequence, and on chromosome 13 many imbalances are likely driven by either epimutations or combinatorial effects of many common variants, which sequence models struggle to predict.

Finally, we visualized examples of well-characterized epigenetically driven allelic imbalance in the autosomes. *LRRTM1* on chromosome 2 is active in EBV-transformed human lymphoblastoid cell lines and exhibits mono-allelic paternal expression. Annotated CpG islands associated with the predicted promoter, coding exon, and upstream were previously found to not be overall differentially methylated [35]. However, MethylSeqNet operating annotation-free from just the sequencing data successfully predicts differential chromatin accessibility between haplotypes (Fig. 3f). The methylation attributions show that the importance of the differential methylation state varies greatly across the window, depending on the sequence context, but the correctly predicted accessibility pattern at the TSS does not cleanly line up with just the methylation landscape. Visualizing four additional examples of imprinting loci in the MethylSeqNet test set establishes that most can be predicted well (Supplementary Fig. 5a-d).

### 2.4 MethylSeqNet conditioned accessibility predictions help interpret rare genetic disease states

MethylSeqNet’s success at predicting allelic imbalance raised the possibility of applying the model to complex rare disease cases where epigenetic signatures can be important indicators or even causal drivers of disease. We applied MethylSeqNet to a long-read sequencing dataset [10] for an Undiagnosed Disease Network patient with a balanced translocation between paternal chr13 and chrX (Methods). The translocation rearranges chrX and chr13 to create der(13) and der(X) (Fig. 4a), while the maternal chromosomes, chr13_*int*_ and chrX_*int*_, remain intact. This patient’s phenotype includes developmental delay and bilateral retinoblastomas. Chromatin accessibility (Fiber-seq) and RNA expression (long-read transcript Iso-seq sequencing) measurements collected in skin fibroblasts and a retinal organoid derived from the patient were used to identify molecular events with potential mechanistic impact on the patient disease state, as summarized in Table 2 from [10]. *NBEA, PDK3, MAB21L1*, and *RB1* are all on either der(X) or der(13), some directly adjacent to the fusion junction (Fig. 4a), and prior work has established clear disruptions in their expression or function. *XIST* and the X-inactivation control locus are on both the intact chrX and the der(X), and their presence on der(X) results in partial X-inactivation of der(X) for a subpopulation of cells.

**Fig. 4.**
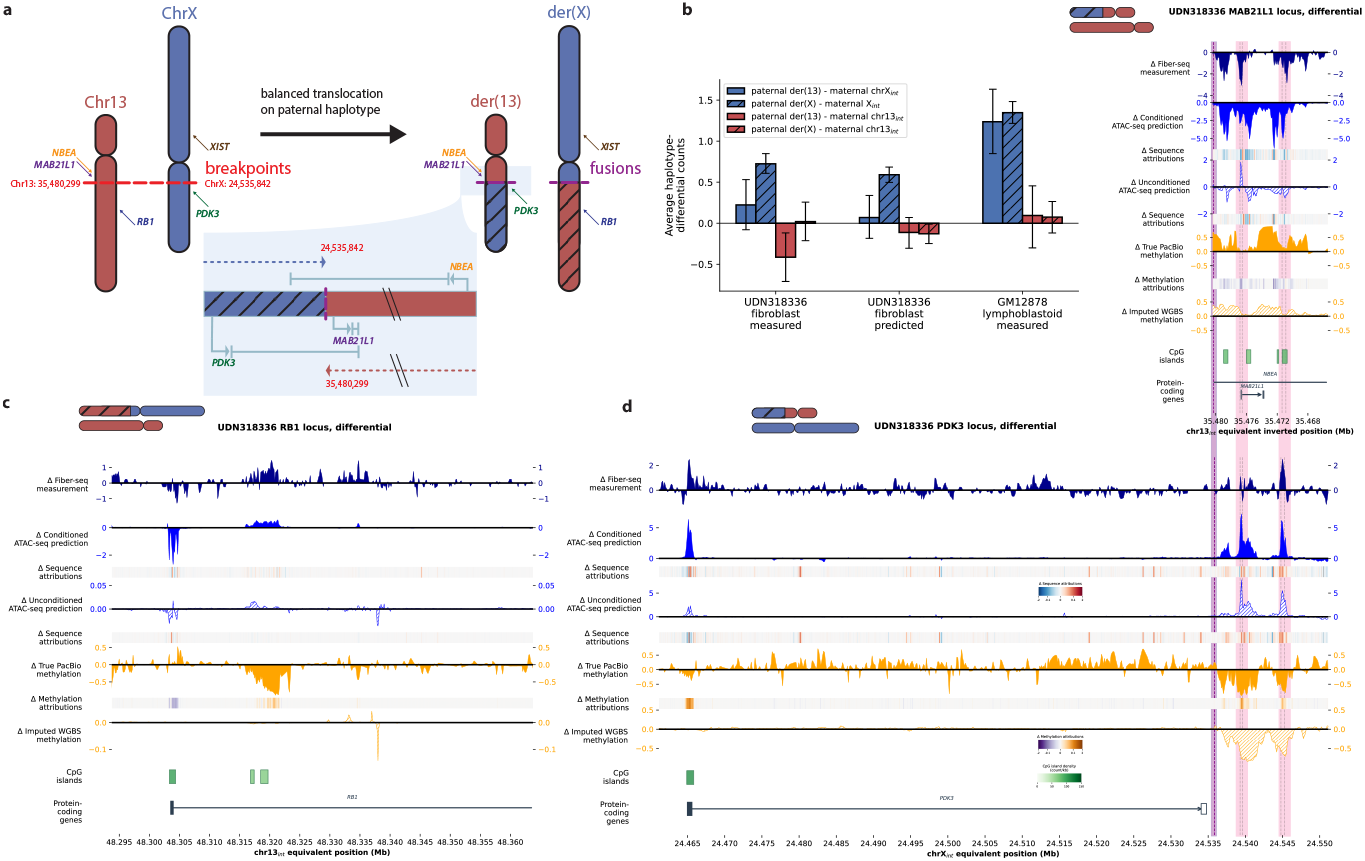
MethylSeqNet predicts functional activity changes characterized in a rare disease patient. **a**, Illustration of the patient studied in [10], where a balanced paternal haplotype translocation creates der(13) and der(X), with several disease-associated genes indicated. **b**, Comparison of the patient’s haplotype-differential accessibility measured in fibroblasts (left) to that predicted by MethylSeqNet for fibroblasts (center). A more typical pattern is shown by measurements for GM12878 in lymphoblastoid cells (right), on which MethylSeqNet was trained. 95% confidence intervals from 100 bootstrap resamples (N = count of genes in chromosome segment, with replacement) are shown. **c**,**d**, Examples of loci where MethylSe-qNet improves haplotype-differential accessibility prediction for retinal organoid: *RB1* experiences partial X-inactivation, *PDK3* activity increases due to adoption of a *MAB21L1*-associated enhancer, and *MAB21L1* is partly silenced by *PDK3* readthrough. We visualize haplotype-differential genomic tracks for accessibility measurements, conditioned and unconditioned ATAC-seq predictions from MethylSeqNet’s neuron task along with their respective sequence attributions calculated using integrated gradients, PacBio methylation measurements, MethylSeqNet imputed WGBS methylation for neurons, methylation attributions calculated using integrated gradients, and CpG island and gene annotations. The fusion junction (purple) and relevant promoters/enhancers (pink) are highlighted.

To investigate whether MethylSeqNet can generalize to predict accessibility changes for this unseen rare disease case, we examined allelic imbalance predictions across loci of interest in both fibroblasts and retinal organoid, comparing with the Fiber-seq measurements. For both cell types, we used model tasks trained on short-read data without allelic imbalance (Methods). First, to examine whether MethylSeqNet can capture large-scale accessibility changes in this setting, we predicted differential accessibility across all TSSs on the intact and fusion-derived chromosomes of the skin fibroblast sample using reference sequence, SNPs, and CpG methylation for sites that do not have the fusion junction in the model receptive field (i.e. within 0.5 Mb). MethylSeqNet successfully predicts weaker chromosome X imbalance between the paternal and maternal X-derived loci compared to GM12878 lymphoblastoid cells, with the regions of chromosome X that become part of der(13) in the paternal copy for UDN318336 having especially weak imbalance (Fig. 4b). Measured imbalance on chr13-derived loci is small on average, although the subset of sites upstream of the fusion site on chromosome 13 shows substantial imbalance only partly captured by MethylSeqNet predictions: the model appears to predict too little imbalance, although the small number of TSS sites on this short region precludes meaningful statistical characterization. Overall, we observe good evidence of generalization to the sequence variants and unusual epigenetic profile for this patient.

Next, we examined specific loci described as disease-associated in [10]. Of these, increased activity of *PDK3* due to enhancer adoption on der(13), partial X-inactivation of *RB1* on der(X), and transcriptional readthrough silencing of *MAB21L1* on der(13) are observed with readouts for which MethylSeqNet has corresponding prediction tasks. *MAB21L1* itself is thought to be unrelated to the disease phenotype. Nonetheless, since the maternal haplotype remains intact, an enhancer associated with *MAB21L1* remains active enough to potentially drive the observed ectopic expression of *PDK3* and the *PDK3-MAB21L1* fusion kinase transcript. With sufficient data, training the model to predict transcript isoforms from sequence could potentially enable direct predictions of fusion transcripts conditioned on the measured epigenetic state, unlocking a key element of the disease-associated molecular phenotype of *NBEA* and *PDK3-MAB21L1*. However, this is outside the scope of the current study.

MethylSeqNet successfully predicts *RB1* silencing as a result of partial X-inactivation of der(X) (Fig. 4c) because there is clear differential CpG methylation, despite the locus existing 13 Mb away from the fusion junction. This demonstrates that the model can generalize to predict inactivation on a chromosome that never exhibits this sort of inactivation in training.

*MAB21L1*, less than 4 kb from the fusion junction, has a different proposed mechanism of silencing: *PDK3* transcripts extend across the fusion junction and through the *MAB21L1* promoter, resulting in readthrough silencing. Because readthrough silencing creates epigenetic shifts [36], MethylSeqNet can predict it through CpG methylation conditioning (Fig. 4d, top). MethylSeqNet also captures that the *MAB21L1* enhancer-like element downstream of the gene body remains accessible in the paternal haplotype, just at a reduced activity level (Supplementary Fig. 6a,b). The unconditioned sequence model predicts allelic imbalance in activity but incorrectly predicts elevated activity at the *MAB21L1* TSS. We demonstrate that the unconditioned prediction is driven primarily by the fusion junction: local SNPs alone drive weaker differential activity, but in both cases the predicted differential pattern from unconditioned sequence aligns poorly with the measurement data (Supplementary Fig. 6c, upper).

For the paternal copy of *PDK3*, 70 kb from the fusion junction, the remaining activity of the *MAB21L1* enhancer-like element on der(13) results in a new distal signal from across the fusion junction. The new enhancer is weakly detected by the unconditioned sequence model, reflected in paternally biased activity for *PDK3* (Fig. 4d, bottom). Without the fusion sequence, the unconditioned sequence model predicts a smaller activity shift with inconsistent sign (Supplementary Fig. 6c, bottom). MethylSeqNet, however, correctly predicts much stronger paternal bias by leveraging the local differential methylation (Fig. 4d, bottom). This result supports our hypothesis that MethylSeqNet effectively leverages local epigenetic signatures to detect the distal regulatory interactions often missed by traditional sequence models.

### 2.5 MethylSeqNet activity predictions are driven by known methylation-sensitive transcription factor motifs

We performed a systematic in silico motif insertion analysis and quantified learned motif effects on activity, conditioned on different methylation landscapes, to investigate whether MethylSeqNet has learned methylation-dependent regulatory grammar. Synthetic motif insertion, inspired by previous work [37], enabled us to explore a larger combinatorial space with higher statistical power than is available with endogenous sequence. Specifically, we designed an in silico analysis to interrogate TF activity under different methylation conditions for 301 specific TFs. To determine methylation levels, we noted that the accessible peak methylation fraction distribution for WGBS when interpolated between CpG sites (Methods) is concentrated around 0.03 and 0.95 (Supplementary Fig. 7a), which we interpret as active and repressive epigenetic states, respectively. We quantified predicted effects for both lower/active (0.03 around the motif) and higher/repressive (0.95 across the input) methylation landscapes across all cell types (Fig. 5a, Methods).

**Fig. 5.**
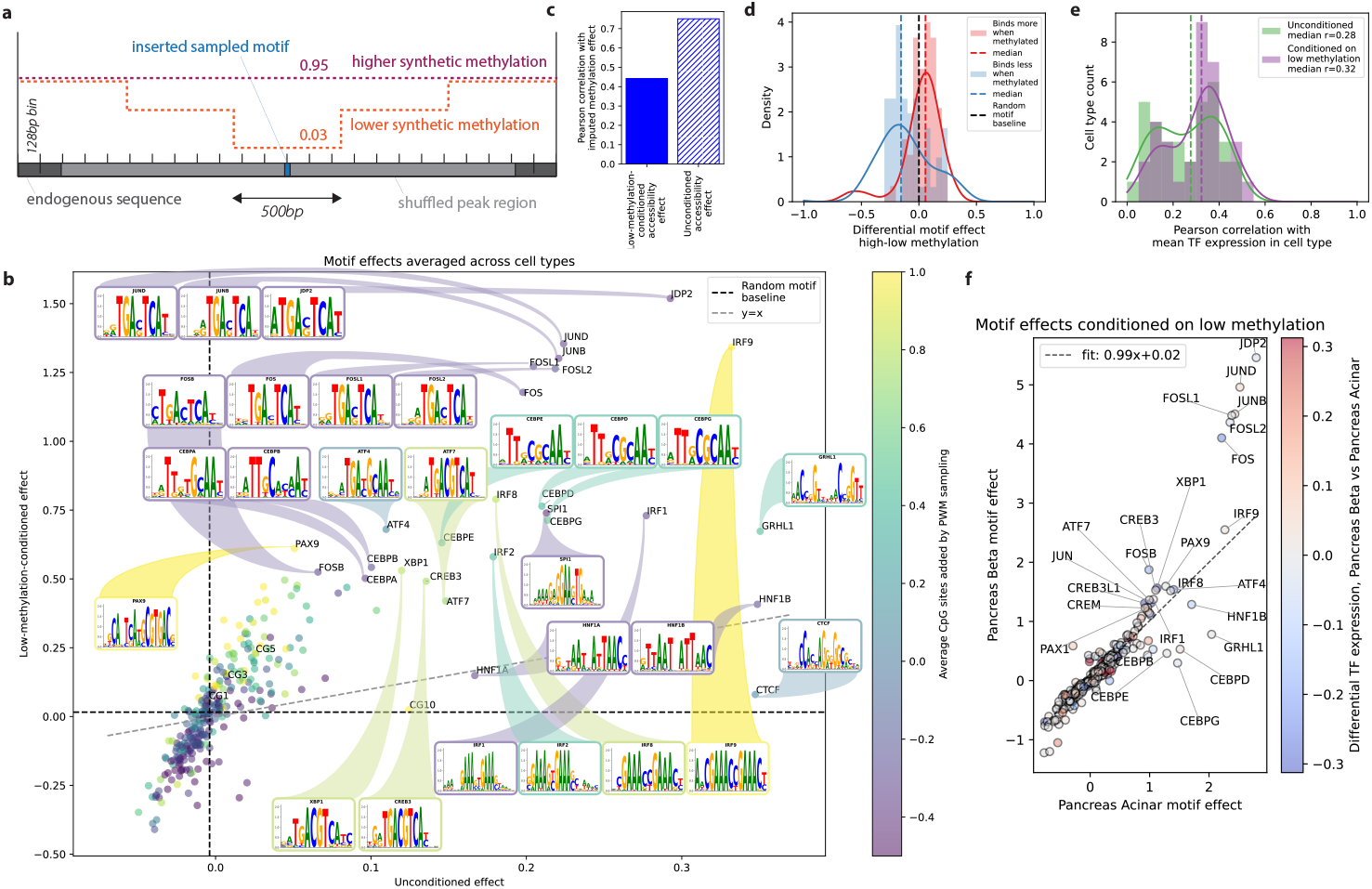
In silico motif insertion analysis of 301 TFs reveals learned sequence-methylation interactions. **a**, Schematic of our motif insertion method, illustrating how a sampled TF motif is inserted at the center of a shuffled peak region under both high and low methylation landscapes. **b**, Scatterplot of low-methylation-conditioned TF motif effect against unconditioned TF motif effect for all 301 TFs studied, averaged across all cell types. Points are colored by the average number of CpG sites added by a sampled position-weight matrix for that TF (the scale is truncated to exclude the all-CpG controls CG10, CG5, etc.), and several of the highest-effect TFs are highlighted. Black horizontal and vertical dashed lines indicate the calculated motif effect of a uniform random motif of length 10 for low methylation and unconditioned cases, and the gray dashed line indicates y=x. **c**, Correlation of MethylSeqNet’s imputed methylation motif effect with accessibility effect conditioned on low methylation (left, solid) and no methylation conditioning (right, striped). **d**, Histograms of differential motif effects (calculated as high methylation effect minus low methylation effect) for TFs known to bind more with methylation (red) vs those known to bind less (blue). **e**, Histograms of Pearson correlations per cell type between chromatin accessibility TF motif effect and corresponding TF expression in each cell type, plotted for low-methylation-conditioned vs unconditioned predictions. **f**, Scatterplot comparison of the low-methylation-conditioned TF motif effects calculated between pancreatic beta and acinar cells, colored by the TF’s difference in expression between the two cell types.

To quantitatively assess which TF motifs are most methylation-dependent in MethylSeqNet’s predictions, we defined motif effect as the predicted mean activity of inserting each motif in the low methylation landscape, averaged across peaks and cell types (Methods). We focused on accessibility motif effect, but also considered expression and imputed methylation effects. Since MethylSeqNet’s predictions reflect the biological fact that high methylation often represses activity, we reasoned that the lower methylation landscape would be more informative for disentangling the relative methylation-conditioned activity of individual TFs. Comparing low-methylation-conditioned accessibility motif effects against those from an unconditioned sequence model, we generally observed a strong relationship between conditional and unconditional predictions, with lower-effect TFs having a roughly linear relationship and higher-effect TFs fanning out into those showing a relatively stronger unconditioned effect or relatively stronger conditioned effect (Fig. 5b). The average slope of the scatterplot’s best fit line is substantially greater than one: conditioning on lower methylation often shifts the model towards predicting high activity for motifs that are recognized from the training corpus.

Many of the TFs with highest predicted effects (labeled in Fig. 5b) match known biology. For example, bZIP family members, including AP-1 complex components (Jun, Fos, ATF, and MAF subfamilies) and the C/EBP family exhibit strong low-methylation-conditional effects consistent with in vitro evidence that bZIP TFs are among the most methylation-sensitive TF families [38]. Conversely, TFs exhibiting strong sequence-only effects and smaller low-methylation-conditional effects include the architectural protein CTCF, which is ubiquitously expressed and drives low-methylation states at tens of thousands of sites genome-wide [28, 39, 40], and endoderm lineage-specifying factors HNF1A and HNF1B. This observation recapitulates the known distinction between signal-dependent TFs, whose binding is gated by local epigenetic state, and lineage-determining TFs, whose chromatin accessibility is defined primarily through sequence recognition [41]. These high-effect TFs represent a variety of position-weight matrix motifs (Fig. 5b), demonstrating that both the conditioned and unconditioned models have learned complex sequence grammar.

We recapitulated the importance of CpG dinucleotides in motifs in driving methylation sensitivity (Fig. 5b) while also showing that more complex sequence features also remain crucial in the conditional representation (Supplementary Fig. 7b). The model has therefore learned to utilize projections of the rich pretrained sequence representation to capture which elements’ activities are conditional upon CpG methylation, regardless of whether the motif contains a CpG site. The mean of absolute shift across conditional features with motif insertion is itself strongly predictive of mean effect across cell types, suggesting that conditional neural network activations from FaRM may be able to serve as a proxy for the methylation sensitivity of an input sequence as part of interpretability analyses.

To probe the model’s implicit understanding of motif impacts on methylation, we calculated motif effects for imputed methylation predictions and found CTCF to be an outlier with very strong imputed methylation, meaning the presence of its motif is a strong indicator of low methylation for MethylSeqNet (Supplementary Fig. 7c). To further disentangle the relationship between low-methylation-conditioned and unconditioned activity, we examined the correlation between MethylSeqNet’s imputed methylation effects and predicted motif effects for both cases. Unconditional motif effect correlates more strongly with imputed methylation motif effect than does conditional motif effect (Fig. 5c), validating the idea that successful unconditioned sequence-to-activity predictions implicitly predict methylation shifts along the way. In contrast, the conditional model uses the methylation provided by conditioning to infer activity separately from whether the TF itself drives differential methylation.

To assess whether MethylSeqNet’s accessibility motif effects agree with experimental findings, we visualized expression levels for TF genes versus the consensus of several assays for methylation’s directional effect on TF binding [38]. MethylSeqNet meaningfully distinguished between TFs that preferentially bind in unmethylated DNA as opposed to those that preferentially bind methylated DNA (Fig. 5d). Similarly, we find that these motif effects have a modest positive correlation with gene expression for their corresponding TFs, slightly stronger than that of the unconditioned model (Fig. 5e). It is likely that the activity of a given TF, predicted from an unconditioned sequence in a given cell type, depends on local sequence context other than its canonical motif. MethylSeqNet’s conditioning helps inform the model whether chromatin state for a sequence element will promote or repress binding, which in turn may explain why conditioned motif effects better capture the presence of TFs in the *trans* environment.

To explore cell-type-differential motif activity, we inspected the pairwise comparison of accessibility motif effects between two pancreatic cell types, acinar and beta cells, which showed among the largest marginal improvements with MethylSeqNet conditioning. We compared their low-methylation-conditioned motif effects (Fig. 5f). While many motifs share similar effects, a subset shows strong differential activity. For example, AP-1 complex motifs have substantial effects in both cell types but have larger effects in beta cells. AP-1 factors are highly active in beta cells, mediating transcriptional activity for glucose metabolism [42, 43] and cytokine response in type 1 diabetes [44]. Similarly, IRF9 shows higher effect in beta cells, consistent with IRF9’s critical role in IFN-*α* response [45–47]. On the other hand, CEBP family TFs, which are established regulators of terminal differentiation in secretory epithelial cells [48, 49], have higher effects in acinar cells. Pancreatic acinar cells have the highest rate of protein synthesis of any mammalian cell type [50], making CEBP enrichment in acinar over beta cells consistent with exocrine secretory identity.

Finally, we applied the same analyses using expression instead of accessibility and observed similar concordance with known biology in most cases. Low-methylation-conditioned effects are stronger than unconditioned effects, and key activity-driving TFs appear with strong effects (Supplementary Fig. 8a). Unconditioned effect correlates well with imputed methylation effect (Supplementary Fig. 8b) and both low-methylation-conditioned and unconditioned TF effects correlate with TF expression levels (Supplementary Fig. 8c). However, weaker separation of TFs annotated by methylation-binding preference (Supplementary Fig. 8d) indicates that the model has a better understanding of methylation’s impact on accessibility compared to expression. In sum, predicted effects for a wide range of motifs reveal that MethylSeqNet has learned the methylation-dependent regulatory impact of hundreds of TFs. Relationships learned by the model reflect known biological effects, suggesting that the conditioning mechanism successfully maps epigenetic regulatory grammar in addition to the motif grammar typically learned by sequence models.

## 3 Discussion

MethylSeqNet represents a new transfer learning and conditioning paradigm for sequence-to-activity models, anticipating the increasing availability of CpG methylation data provided by long-read DNA sequencing to improve functional activity predictions and enable joint interpretation of sequence and methylation effects. By training jointly on CpG methylation, chromatin accessibility, and gene expression measurements from short-read data for dozens of primary cell types and from haplotype-phased long-read data, MethylSeqNet learns epigenetic influences on regulatory grammar across a wide range of cellular contexts.

Our results suggest that conditioning on methylation state results in greater improvements for chromatin accessibility than for gene expression, and as a result the analysis in this study focuses primarily on chromatin accessibility predictions. The disparity lines up well with known biology: there is a tighter local coupling observed between CpG methylation state and chromatin accessibility as compared to gene expression [51, 52]. CpG methylation at regulatory elements is closely coupled to TF occupancy and local chromatin state [51], whereas its relationship to transcriptional output is more indirect and context-dependent [52]. This is reflected in the methylation atlas data used for cross-cell-type training, in which the methylation variation most relevant to cell-type-specific gene expression is predominantly found at distal enhancers rather than at TSSs themselves [14]. These dynamics makes successful methylation-conditioned expression predictions challenging: like all leading sequence models [7, 53], MethylSeqNet struggles to model distal effects on activity, including methylation effects. We demonstrated that performance boosts from increasing the receptive field for conditioning state drop off beyond 8 kb. We hypothesize that there may be insufficient examples in the training data for any given sequence grammar of long-range interactions in terms of methylation-mediated activity. Our modular conditioning framework should make it possible to apply any future methods that help learn distal interactions from relatively sparse data in sequence models.

Unconditioned sequence models are expected to perform fairly well on cell-type-differential activity predictions because cell lineage and identity are often consistent across healthy individuals and driven by sequence elements recognized by TFs and other molecular machinery. Indeed, MethylSeqNet performance indicates that the methylation conditioning yields only modest improvement on average across the full test set. However, the methylation-conditioned model nevertheless makes dramatically better cross-cell-type predictions at thousands of individual loci, typically by identifying active elements modulated by epigenetic state that the unconditioned sequence model misses. Understanding precisely where and why methylation conditioning improves predictions could inform refined architectures or training frameworks to better capture distal effects or rare sequence element effects.

MethylSeqNet is especially powerful for predicting allelic imbalance of chromatin accessibility, which is often epigenetically driven. As expected, unconditioned sequence models are unable to predict X-inactivation given haplotype-specific SNPs alone, but conditioning on CpG methylation allows MethylSeqNet to predict the strength of differential chromatin accessibility. We also demonstrated that MethylSeqNet predicts substantial allelic imbalance on autosomes with very few imprinting loci, which is missed by unconditioned sequence models. We posit that this observation is driven by two distinct phenomena. First, random silencing of one haplotype is commonly observed for many genes and accessible chromatin sites: while distinct from X-inactivation and imprinting, this phenomenon is maintained by epigenetic signatures that CpG methylation conditioning can account for. Second, sequence variants that drive differential chromatin accessibility and gene expression often also drive differential CpG methylation [15]. While state-of-the-art unconditioned sequence models only partly capture the effects of these variants, CpG methylation conditioning can improve predictions simply by providing a stronger signal for activity that can be difficult to learn from sequence alone.

The largest limitation for MethylSeqNet is that the available methylation training datasets are relatively small. This limitation informed our decision to use a powerful pretrained sequence encoder in the MethylSeqNet architecture. Collecting and matching independently profiled methylation and activity data is labor intensive and limited by the match quality of the different cellular contexts in which measurements were collected. Training on the growing corpus of Fiber-seq datasets from diverse cell types or individual donors offers a promising route forward because the methylation and activity measurements are directly coupled [9, 10]. Another limitation is that measurements from healthy cell types capture only a subset of the relationship between sequence activity and methylation. Training on high-throughput perturbation data, such as methylation-sensitive massively parallel reporter assays [54] could help the model generalize beyond healthy cell types. We anticipate that scaling up such datasets and achieving better signal-to-noise ratios will be critical enablers for using such data to improve methylation-conditioned models. Given limited data, we used in silico motif insertion analyses to demonstrate that MethylSeqNet predictions are often concordant with known biology. While this analysis does not directly validate model performance with real quantitative data, it provides additional evidence that MethylSeqNet’s conditioning mechanism learns biologically relevant signals.

We demonstrate that MethylSeqNet can successfully generalize to large-scale and local activity differences in an undiagnosed disease patient carrying a balanced translocation that produces widespread allele-specific epigenetic shifts. The ability to make such predictions from haplotype sequence and CpG methylation alone—which to our knowledge is novel among sequence-to-activity models—enables disease-associated activity inference from standard long-read whole-genome sequencing datasets, leveraging the CpG methylation signal natively captured by PacBio HiFi or Oxford Nanopore platforms. Clinical variant pathogenicity classification relies on combining independent lines of evidence, among which computational predictors of variant impact are an established and increasingly weighted category [55, 56]. MethylSeqNet extends such computational prediction to non-coding regulatory variant effects conditioned on epigenetic state, complementing experimental functional readouts such as Fiber-seq. Its input attributions can further help disentangle which specific sequence elements or epigenetic states within a region are most likely mediating an observed functional effect, aiding variant prioritization where multiple candidates co-occur.

Future work could apply this analysis to many more rare disease patients. To improve application to relevant disease states, it is likely that training on organoid-like cells will be valuable. The current model struggles with epigenetic signatures characteristic of organoids and embryonic cells, but such systems are a promising way to model the effects of unusual sequence variation in specific tissues. It may also be possible to extend the MethylSeqNet framework to infer methylation levels in primary cell types based on methylation profiles from patient-derived cell lines. Some loci preserve methylation state for a given individual across differentiated cell types while others undergo reprogramming. This relationship could be modeled with appropriate training data, enabling diagnostic assessment of patient’s epigenetic state from easily-collected samples.

There are several other potentially fruitful opportunities for MethylSeqNet that we did not explore in this study. For instance, predicting which of the thousands of methylation changes in a pathogenic cell state are driving which relevant activity differences may help better understand the mechanisms involved and assist in treatment design. T cell exhaustion is a disease-relevant example of programmatic regulatory shifts with a major epigenetic component [57–60]. As we demonstrated, methylation alone is an inadequate predictor of differential activity because of its varying interactions with sequence elements; applying MethylSeqNet to jointly parse the interactions of sequence and methylation may further elucidate biological mechanisms. Predicting epigenetic edit effects is another potential application. As epigenetic editing becomes an increasingly refined and scalable technique, models that can predict the functional effects of locus-specific edits may help select edit sites for a given target gene. A longer-term goal would be to predict the effects of personal genetic variation and potential epigenetic edits in tandem, i.e., to design epigenetic therapies for personalized medicine. Since edited epigenetic states may go well beyond the distribution of endogenous epigenetic states, such extensions remain challenging without significantly more experimental data characterizing this combinatorial space.

To our knowledge, MethylSeqNet is the first epigenetically conditioned sequence-to-activity model, and we anticipate a wider range of future applications than those mentioned here. We have open-sourced the MethylSeqNet model and the core computational libraries that implement it. While we demonstrate the framework on supervised CNN and transformer models, the concept can extend to any sequence model whose embeddings contain information about gene regulation, including DNA language models [61–64]. Furthermore, the transfer learning framework can be easily adapted to condition activity predictions on histone state, condition expression predictions upon chromatin accessibility, or even condition on information about the *trans* environment, such as TF abundance or metabolic state. By making this principled and practical epigenetic-conditioning framework for sequence modeling available to the broader research community, we hope to help accelerate disease diagnostics and increase our understanding of epigenetic effects in the regulatory genome.

## Methods

### Cell types preprocessing

We reference three different primary cell type atlas datasets in our analysis. Cell-sorted bulk bisulfite sequencing from [14] provides CpG methylation genome-wide for 39 cell types. Single-cell ATAC-seq from [23] provides chromatin accessibility for 222 cell types from 30 tissues. Finally, the Fantom5 consortium [24] provides cell-sorted CAGE-seq for TSS activity. Methylation inputs were matched to functional activity outputs as described in Supplementary Table 1.

For the methylation atlas, we downloaded processed bigwig files with dinucleotide-level CpG methylation fractions genome-wide from GEO accession GSE186458. These were processed as described in [14]. For the chromatin accessibility atlas, we downloaded bigwig files with processed counts-per-billion aligned counts from https://decoder-genetics.wustl.edu/ catlasv1 (raw data available under GEO accession GSE184462). These were processed to pseudobulk bigwig tracks as described in [23]. For the TSS activity atlas, we downloaded TSS counts bed files from FANTOM 5 https://fantom.gsc.riken.jp/5/datafiles/reprocessed/hg38_v9/ for human primary cells [24]. We downloaded the latest reprocessed files from the FANTOM5 consortium, updated in 2021. All downloaded files were aligned to the hg38 reference genome.

We preprocessed the methylation data by creating a “CpG-methylation-possible” mask for all sites with a valid CpG methylation fraction measurement and extracting the 0-to-1 fraction methylated at each of these sites. This was then combined with the one-hot DNA sequence encoding to create the input tensors.

We preprocessed the accessibility and expression label data by normalizing all counts to counts-per-billion genome-wide, binning into 128-bp bins, and taking the square root of normalized counts exceeding 32 for ATAC-seq and 384 for CAGE-seq. This preprocessing was designed to be similar to Basenji2 and Enformer [3, 5, 6] which are optimized for accessibility and TSS activity predictions.

### Allelic imbalance preprocessing

Allelic imbalance data, collected using PacBio Iso-seq and Fiber-seq, was sourced from and processed as described in [10]. We used data from a GM12878 lymphoblastoid cell line for training and data from UDN318336 skin fibroblast and retinal organoid, including a haplotype-phased genome assembly, to assess generalization to rare disease interpretation. Processed GM12878 data is available for download from https://s3-us-west-1.amazonaws.com/stergachis-manuscript-data/2023/Vollger_et_al_long-read_multi-ome, while UDN data access is through the UDN dbGaP (Access instructions: https://sharing.nih.gov/accessing-data/accessing-genomic-data/how-to-request-and-access-datasets-from-dbgap).

Haplotype-phased Fiber-seq accessible counts data were preprocessed by running counts-per-billion normalization in 128 bp bins and taking the square root of residual counts exceeding 32.

For Iso-seq phasing and constructing personalized genome fasta files, we used a precomputed phased vcf file downloaded from UCSC Platinum Genomes for GM12878 and created a phased vcf for UDN318336 using the haplotype-phased assemblies using `run-dipcall` with `dipcall v0.3`. Using the phased vcf files and the hg38 reference genome, we used `bcftools consensus -i ‘TYPE=“snp”‘` with `bcftools v1.21` to create fasta files for each haplotype with a consistent coordinate system. To phase the Iso-seq coverage to maternal and paternal haplotypes, we first used `samtools index` from `htslib v1.21` where needed to index bam files then used `whatshap haplotag` with `whatshap v2.4` with the phased vcf files. Then, to process the Iso-seq coverage to be comparable to the CAGE-seq TSS activity, we used `isoseq collapse` with `isoseq v4.3.0` to map the 5’ and 3’ ends of transcripts, applying `--min-aln-coverage 0.85` to avoid filtering out an overly large fraction of reads. The `isoseq collapse` output .gff and .abundance.txt were used to count transcript 5’ ends in each 128 bp bin. To correct for losses due to haplotype phasing, counts-per-billion 5’ end abundance was calculated with unphased Iso-seq reads and then this value was used to normalize phased counts in each bin, such that total phased counts in each bin were equal to total unphased counts-per-billion. Bin counts were softclipped by taking the square root of residual counts exceeding 384, similar to [5].

### ConditionedSeqNN framework

MethylSeqNet is implemented within the ConditionedSeqNN Python framework, built on PyTorch Lightning. DNA sequence and methylation state pass into a sequence encoder and conditioning state encoder. The embeddings from the sequence encoder and a representation of the methylation conditioning state are passed into a conditioning mechanism, the output of which passes into an output head to make the multitask predictions.

The sequence encoder is implemented as a `pytorch nn.ModuleList` that runs on one-hot encoded sequence. For the results presented in the main text, the conditioned and unconditioned models were trained using Borzoi replicate 0 from https://github.com/johahi/borzoi-pytorch, for which **h** has 1920 channels. Train-from-scratch experiments and the associated pretrained comparisons of Basenji2 used https://github.com/d-laub/basenji2-pytorch, for which **h** has 1536 channels. To maintain easy compatibility between the Basenji2-based, Borzoi-based, and small CNN experiment models described below, we chose a 128 bp bin size for labels and predictions. Compared to Borzoi’s RNA coverage tasks [6], our tasks did not require this finer resolution. Thus, for Borzoi embeddings, we downsampled embeddings by a factor of 4.

The conditioning state representation is calculated from the 0-to-1 fraction methylated per CpG site. First, methylation fraction is linearly interpolated between CpG sites to create a more stable representation regardless of CpG density. Second, the interpolated signal is averaged in 128 bp bins to align with the downsampled sequence embeddings and labels, resulting in one scalar CpG methylation representation per positional bin.

Our proposed conditioning mechanism, Factorized Representation Modulation, described in Results 2.1, is optimized for the methylation conditioning case with many methylation states (e.g. different cell type methylation landscapes) for the same sequence input. However, we also baseline against the similarly expressive Feature-wise Linear Modulation (FiLM) [65]. The FiLM equation (Eq. 3), which can be contrasted with FaRM (Eq. 2), is

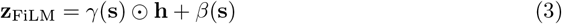

where *γ* and *β* are learned functions of the conditioning state **s**. Because many low-dimensional conditioning states (cell types) are applied to the same sequence, standard FiLM overparameterizes the modulation by learning 2d modulation coefficients as a function of s. FiLM is therefore slower to run for similar predictive performance, whereas FaRM is nearly as efficient as simply concatenating scalar methylation state to the pretrained embeddings while performing substantially better.

Finally, the output head integrates spatial information from the new conditioned representation **z**. We implement the output head as a dilated residual CNN as described in [3], preceded by a fully connected layer to process the conditioned representation to appropriate dimension and followed by a fully connected layer and softplus activation to calculate final predictions from the CNN outputs. The model described in Results uses 5 dilated convolution blocks with a kernel size of 3 and dilation rates of 1, 2, 4, 8, and 16, for a total receptive field of 63 bins, corresponding to an 8064 bp receptive field.

### Training

We trained the model with the same train-test split as Borzoi and a similar multi-task framework. Pretrained sequence encoder parameters were frozen and only the conditioning mechanism and output head were trained. A single output head predicts all tasks, thus sharing parameters except for in the final layer which maps cell-type-specific grammar. This parameter sharing is designed to aid with generalization. However, because different tasks are associated with different conditioning states, the output head is run multiple times per genomic sequence in training. Furthermore, we applied loss to only the subset of tasks for which a given sequence has conditioning methylation inputs. When training jointly on cell types and allelic imbalance data, each sample was randomly selected from one or the other dataset during training.

We used a Poisson loss function for predicted counts, binary cross-entropy for imputed methylation fraction, and the Adam optimizer with learning rate 0.002, betas (0.9,0.999), and 32 samples per optimizer step. The cell types samples take more GPU memory than the allelic imbalance samples because there are more conditioning states to run; backpropagation for cell types samples requires 5.7 GB and allelic imbalance samples require 4.8 GB. All experiments were trained until total validation loss plateaued, typically about 5,000 optimizer steps or about 50 hours on a single A40 GPU. Supplementary Table 1 describes the input-to-output mapping, including task structure and representative performance metrics per task.

### Small CNN experiments

The small CNN models implemented as a sequence+methylation encoder to simplify basic ablation experiments each contained nine convolution blocks. The first takes in the encoded sequence and/or methylation with a filter size of 15, the subsequent 6 have filter size 5 and pool by a factor of 2 after each, the penultimate block has filter size 5 without pooling, and the final block has filter size 1 and filters equal to the number of tasks, effectively serving as a fully connected layer per-position. All prior blocks have 50 filters for a total parameter count of 97 thousand. Models were trained on the full cell types and allelic imbalance datasets. Input data ablation was achieved by trimming the channels coming into the model to only sequence, only interpolated methylation, or both. Methylation smoothing was achieved by applying an all-ones convolution filter to degrade information content in the input signal.

### Factorized model hyperparameter experiments

To establish that MethylSeqNet can jointly model cell type and allelic imbalance data, we performed an ablation training on only one or only the other (Supplementary Fig. 9a). We saw a modest performance reduction from joint training for cell types performance which we hypothesize is due to the model being forced to more strongly factorize the representation (Supplementary Fig. 2e) as discussed in Results. Allelic imbalance performance is not significantly altered by joint training.

To establish that pretraining helps performance, we used the smaller Basenji2 model to enable full training on a reasonable timescale. MethylSeqNet with a pretrained Basenji2 converges much faster and performs better across all cell type tasks, compared to a randomly-initialized Borzoi trained from scratch with the same conditioning (Supplementary Fig. 9b).

The conditioning and output head complexity were selected for layer count and receptive field (Supplementary Fig. 9c), conditional to unconditional features ratio (Supplementary Fig. 9d), and total feature count (Supplementary Fig. 9e). Increasing the receptive field for methylation processing by adding more convolution layers or implementing transformer layers similar to those in [6] showed substantially less improvement after 5 layers for the CNN. 7 layers is slightly better for some tasks but we opted for 5 layers as a more computationally lightweight design. Varying the ratio of conditional to unconditional features for FaRM, keeping the total feature count fixed at 1024 across the two representations, shows relatively small performance shifts between a 1:15 ratio and a 15:1 ratio. The cell types tasks perform best with more unconditional features and the allelic imbalance tasks perform best with more conditional features. In order to balance these, we chose to use a 1:1 conditioned:unconditioned ratio. The extra parameters for the FaRM output head improve performance for most tasks and metrics over the single-layer output head found in the original Borzoi model (Supplementary Fig. 2d). Finally, experiments for total representation features in a balanced 1:1 conditioned:unconditioned FaRM model demonstrate relatively little sensitivity ranging from 256 to 2048, but with some benefit from increasing to 1024 and relatively little benefit from increasing to 2048. Thus we select 1024 features for the presented model architecture.

We trained a version of this optimized architecture for each Borzoi replicate (Supplementary Fig. 9f). As seen elsewhere, Iso-seq task results are the most inconsistent, whereas non-differential accessibility metrics are highly consistent. Cell-type or haplotype differential metrics are less consistent.

### Cell types analysis

We evaluated model performance on held-out test set genomic regions for ATAC-seq and CAGE-seq tasks. Subjective cell-type matching quality, from 0 to 5, and bisulfite sequence read depth were recorded for each WGBS:ATAC-seq and WGBS:CAGE-seq pair. For each cell type, we calculated both a cross-locus Pearson correlation for the raw predictions and targets and a cell-type-differential cross-locus Pearson correlation for mean-subtracted predictions and targets (Eq. 4), defined as

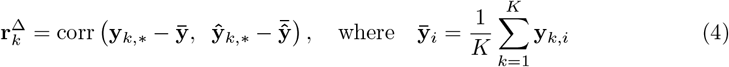

where *k* is the cell type and *i* is the locus. These Pearson correlations were compared between a conditioned and an unconditioned model to show which cell types change the most.

Next, loci were selected to identify where improved performance arises. To focus evaluation on informative loci, we selected a set 𝒮 of peak bins where the maximum count across cell types exceeded a threshold (*thresh*_ATAC_ = 15 processed counts, *thresh*_CAGE_ = 40 processed counts), taking the highest bin within each contiguous above-threshold run and enforcing a minimum spacing of 8 bins (1,024 bp) between retained peaks. We calculated per-locus cross-cell-type Pearson correlation at peak loci (Eq. 5).

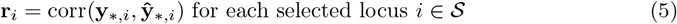

Observing the distribution of locus performance, with most loci having little change from conditioning while some were very substantially improved, we plotted the true measured counts versus conditioned minus unconditioned predicted counts at each locus with per-locus cross-cell-type Pearson correlation improved by more than 0.5. Since some sites have higher counts, i.e. more activity, with conditioning while others have fewer counts, we visualized examples selected from the set of loci for which Δ**r**_*i*_ *>* 0.5 where the conditioned model is closer to the targets by at least 1 count and the log-fold difference between the conditioned and unconditioned prediction exceeds 1. *CEL* is one such example, highlighted in the main text because of its relatively complete annotations. *CEL* is primarily active in pancreatic acinar cells and lactating mammary glands [29]. Mammary luminal epithelial cells are included in MethylSeqNet training, but if the individual donors were not lactating then only the pancreas acinar cells would show strong *CEL* activity. Other examples highlight representative dynamics seen with the selection criteria.

To assess whether chromatin compartments explain the conditioning improvements, we cross-referenced with chromatin compartment data from [31] for cell types shared between their annotated cell types and those used in MethylSeqNet training.

### Allelic imbalance analysis

We evaluated model performance on held-out test set genomic regions for Fiber-seq and Isoseq tasks. Haplotype-differential Pearson correlation is calculated by comparing true paternal - maternal activity with predicted paternal - maternal activity (Eq. 6).

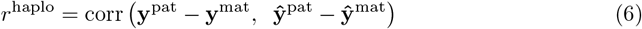

TSSs for chrX and chr13 were identified in hg38 by referencing Gencode basic annotations release 49 (GRCh38.p14) and identifying only canonical transcripts. Reference sequence with SNP variants was processed as described in Allelic imbalance preprocessing, allowing us to run both unconditioned sequence-only predictions and methylation-conditioned predictions and compare with measured haplotype-differential targets.

Too few annotated imprinted genes were in the test set, based on the list provided by https://geneimprint.com/site/genes-by-species, to provide meaningful statistics. However, visualizing several examples revealed that many have differential activity accurately predicted by MethylSeqNet.

### Input attributions

We applied Integrated Gradients [66] using Captum [67], to achieve GPU-accelerated gradient-based attributions. For sequence, a dinucleotide-frequency-preserving shuffle provided a reasonable baseline, with 20 baselines averaged to provide a clean signal. However, the dinucleotide shuffle disrupted CpG positions, making the integral between the baseline and the input out-of-distribution along its entire path. Thus, methylation attributions calculated simultaneously with dinucleotide-shuffle sequence attributions created noisy results with attribution spikes in places that were likely spurious.

Instead, we held methylation roughly constant for the dinucleotide shuffle, interpolating existing methylation landscape to new CpG locations and only recording the sequence attributions. Then, holding sequence constant, we applied a uniform methylation baseline of 0.95, representing a repressed chromatin state for most sequence elements. This produced clean methylation attributions that remained highly interpretable, demonstrating the importance of each input CpG site assuming sequence remains unchanged.

### Genomic track plots

To visualize predictions for an individual locus, we developed a plotting pipeline with many parallel elements. In addition to measured activity and methylation, unconditioned predictions for activity and methylation, and methylation-conditioned activity predictions, we visualized sequence attributions, methylation attributions, gene annotations, high-scoring TF motifs, and CpG islands.

Basic gene annotations were downloaded from Gencode release 49 and used to identify TSSs with associated gene names and categories. This .gtf file was sorted, bgzipped, and tabix-indexed with `hts v1.21`, then queried with `pysam` and subset to protein-coding transcripts. Genome-wide motif matches were downloaded from Non-redundant archetype motif clusters (version 1.0) [68]. CpG islands were generated with `ucsc-twobittofa v469` and `ucsc-maskoutfa v472` using the following command sequence provided by the UCSC genome browser.

~~~
twoBitToFa assembly.2bit stdout | maskOutFa stdin hard stdout \
 | cpg_lh /dev/stdin 2> cpg_lh.err \
  | awk ‘{$2 = $2 - 1; width = $3 - $2; \
  printf(“%s\t%d\t%s\t%s %s\t%s\t%s\t%0.0f\t%0.1f\t%s\t%s\n”, \
  $1, $2, $3, $5, $6, width, $6, width*$7*0.01, \
  100.0*2*$6/width, $7, $9);}’ \
    | sort -k1,1 -k2,2n > cpgIsland.bed
~~~

### Undiagnosed disease patient analysis

Because the UDN318336-derived cells from the study are not directly represented in our training set, we selected the best-fit cell types from training. For skin fibroblasts we selected the primary cardiac fibroblast prediction task and for the retinal organoid, which is fundamentally a neural tissue, we selected the neuron task.

To create fusion sequences adequately capturing the balanced translocation while enabling easy integration with our pipelines designed for reference genome coordinates, we created new fasta and bigwig files containing new contigs derX_*fwd*_, derX_*rev*_, der13_*fwd*_, and der13_*rev*_. Because the translocation fuses the pre-junction-site part (by reference coordinates) of chr13/chrX into der13 and the post-junction-site part of each into derX, the der13 has the X-derived sequence reverse complemented and derX has the 13-derived sequence reverse complemented. By creating the fwd and rev contigs we can run inference on any gene in the 5-to-3 prime direction of its native chromosome, even if it is flipped on the derived chromosome.

Because the sequence fasta, methylation bigwig, and Fiber-seq activity bigwig all contain the fusion contigs, we visualized activity through the fusion junction, as in Fig. 4d. For *PDK3*, the allelic imbalance upstream of the junction contains X-chromosome-native sequence for both alleles, but downstream of the junction we are comparing X-chromosome-native sequence on the maternal copy with chromosome-13-native sequence on the paternal copy. This is precisely what the *PDK3* promoter is exposed to in terms of differential regulatory signals.

### In silico motif insertion analysis

To define a set of motifs, we considered TFs whose binding activity was explicitly characterized with respect to methylation [38], known pioneer TFs [69], and CTCF as a known methylation-sensitive TF [28] for our targeted investigations. We utilized characterized TF motif position-weight matrices from [70]. We also included position-weight matrices representing differing lengths of uniform random motifs and multi-CpG motifs as controls.

To create sequences for inference, we first identified peaks active in at least one cell type. Since endogenous accessibility peaks vary in magnitude and CpG density across cell types, we selected them from randomized cell types. For each peak and TF, we centered the model’s sequence context at each peak, shuffled the central 2 kb while preserving dinucleotide frequencies, sampled a motif from the position weight matrix corresponding to the TF, and centered the sampled motif in the sequence passed to MethylSeqNet. We ran 5 trials per peak×motif×methylation level for high/repressive methylation (0.95), low/activating methylation (0.03), and unconditioned. The predicted mean effect for chromatin accessibility, gene expression, or imputed methylation predictions was calculated separately for each of the 301 TF motifs, averaged across 5 trials in each of 500 peaks for all cell types. We subtracted the shuffled peak sequence to achieve a differential effect.

To facilitate faster analysis with much smaller intermediate output files, thus enabling many more replicates per TF, we truncated the receptive field of our Borzoi pretrained encoder to 16 kb by removing its final cropping layer. Because the effects are always differential relative to a shuffled baseline and the only input change is in the central bin, we posit that the truncated receptive field does not directionally impact results.

## Data availability

Public data used to train MethylSeqNet is available at GEO accession GSE186458 (WGBS data), GEO accession GSE184462 (pseudobulked scATAC-seq data), the FANTOM5 database https://fantom.gsc.riken.jp/5/ (CAGE-seq data), and NCBI BioProject accession PRJNA1124997 (GM12878 Fiber-seq and Iso-seq data). UDN data access is through the UDN dbGaP. Detailed instructions for submitting a data access request can be found on the NIH Scientific Data Sharing website (https://sharing.nih.gov/accessing-data/accessing-genomic-data/how-to-request-and-access-datasets-from-dbgap).

## Code availability

The implementation of MethylSeqNet and the ConditionedSeqNN framework is available at https://github.com/OberonDixon/methylseq-net. Code to reproduce training runs and figure generation is available at https://github.com/OberonDixon/methylseq-net-reproducibility. Trained model checkpoints are available on Zenodo at https://doi.org/10.5281/zenodo.20275120.

## Acknowledgements

We thank members of the Streets and Ioannidis labs for useful discussions and insights, especially Michal Rozenwald, Daniel Lewinsohn, Jeremy Marcus, and Yizi Mao. This research used the Savio computational cluster resource provided by the Berkeley Research Computing program at the University of California, Berkeley. O.D. was supported by the Natural Sciences and Engineering Research Council of Canada (NSERC) Postgraduate Scholarship – Doctoral (PGS D) award number 587658. Research reported in this publication was supported by the National Human Genome Research Institute of the National Institutes of Health under award number R01HG012383 to A.S. A.S. holds the Lester John and Lynne Dewar Lloyd Distinguished Chair in Bioengineering and is supported by the Harvey and Leslie Wagner Foundation. This work was supported, in part, by US National Institutes of Health (NIH) grants 1DP5OD029630 and 1U01HG013744 to A.B.S. M.R.V. was supported by a training grant (T32) from the NIH (2T32GM007454-46). M.R.V. was also supported by a Pathway to Independence award from the National Institute of General Medical Sciences (1K99GM155552-01 and 4R00GM155552). A.S. and N.M.I. are Biohub San Francisco Investigators.

## Author contributions

O.D., A.B., A.S., and N.M.I. conceptualized the factorized model. O.D. implemented the model framework and real data analyses with assistance from A.B. A.B. implemented the motif insertion analysis with assistance from O.D. O.D., A.B., A.S., and N.M.I. interpreted the results. M.R.V. and A.B.S. provided rare disease patient data, interpretation, and clinical pipeline information. O.D., A.B., A.S., and N.M.I. drafted the manuscript. All authors read and approved the final manuscript.

## Supplementary Material

**Supplementary Fig. 1.**
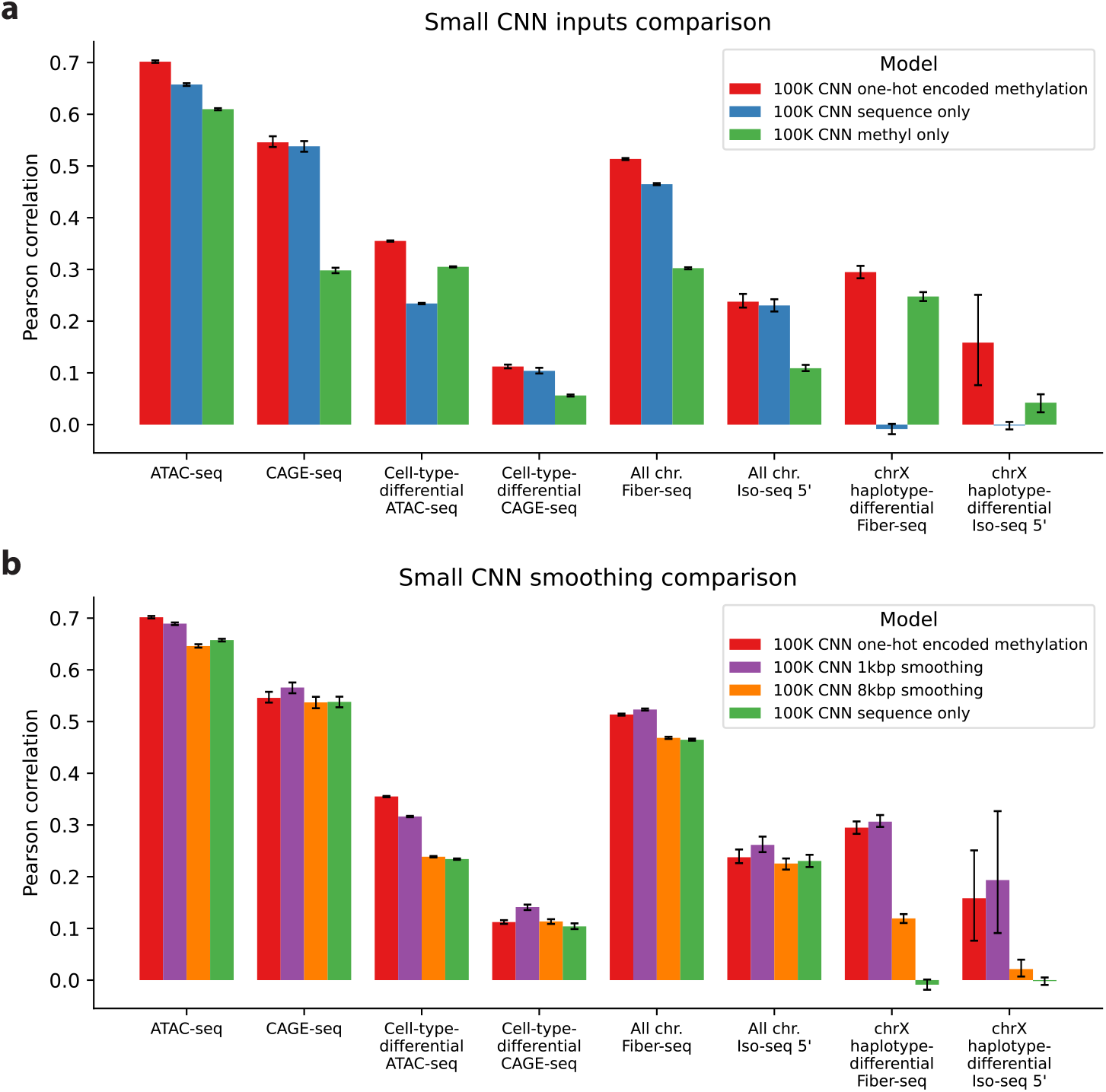
Small CNN encoder ablations. **a**, Performance comparison of small CNN models with sequence and methylation input, just sequence, and just methylation. 95% confidence intervals from 100 bootstrap resamples (N = full test set subset size, with replacement) are shown. **b**, As in a but for small CNN models with different levels of input methylation smoothing.

**Supplementary Fig. 2.**
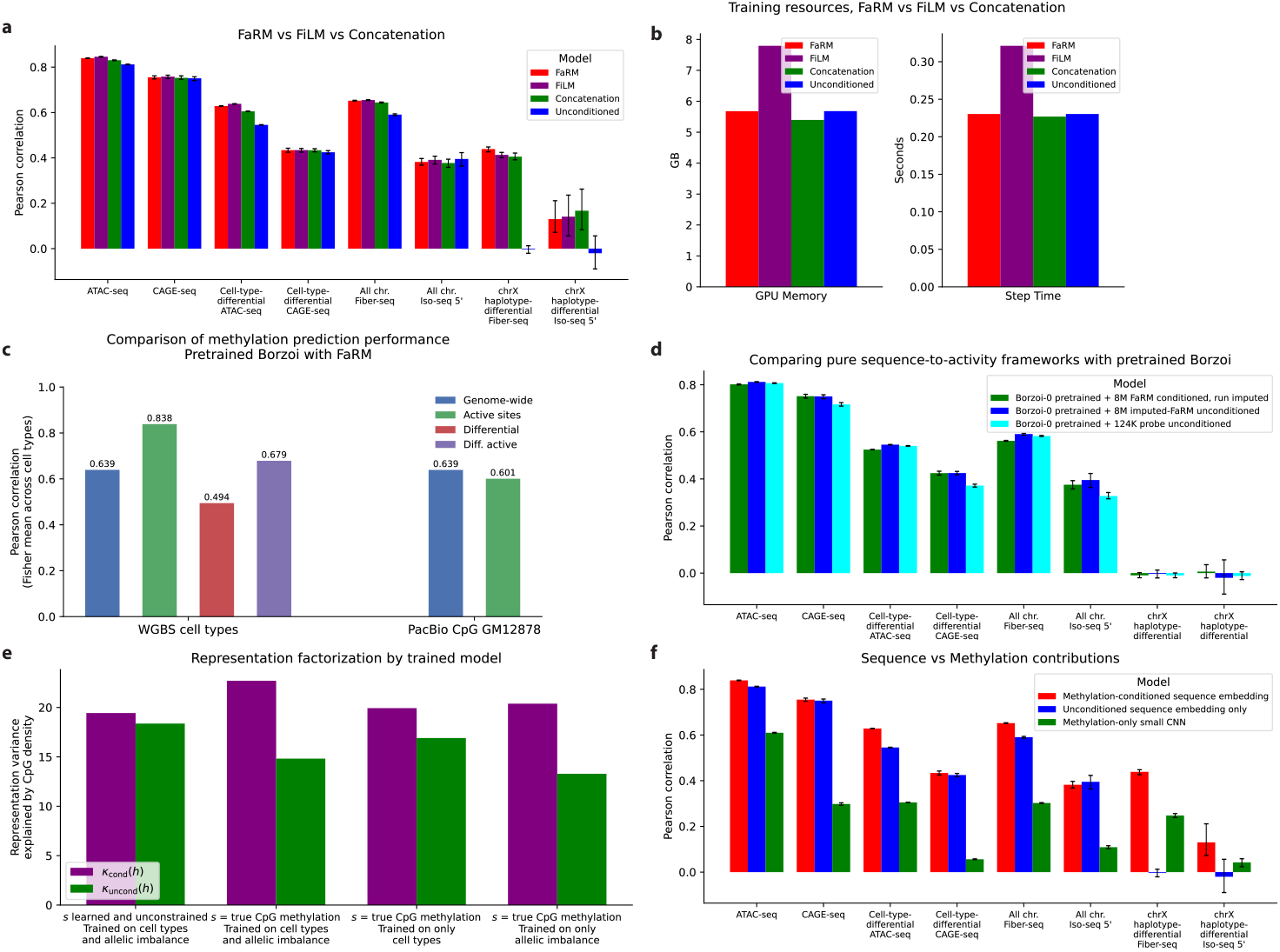
FaRM performance extended. **a**, Prediction performance comparisons of FaRM conditioning, FiLM conditioning, simple concatenation of the conditioning representation, and the unconditioned MethylSeqNet. 95% confidence intervals from 100 bootstrap resamples (N = full test set subset size, with replacement) are shown. **b**, GPU memory and training step time comparisons of FaRM conditioning, FiLM conditioning, simple concatenation of the conditioning vector, and the unconditioned MethylSeqNet. **c**, Imputed methylation performance comparison across cell types and data types. **d**, As in **a** but comparing FaRM output head run with imputed conditioning, an unconditioned FaRM output head, and a linear probe on the Borzoi embeddings. **e**, Comparison of representation variance explained by CpG density for conditional and unconditional representations across different MethylSeqNet configurations. **f**, As in **a** but comparing MethylSeqNet with pretrained Borzoi, both conditioned and unconditioned, and a methylation-only CNN model.

**Supplementary Fig. 3.**
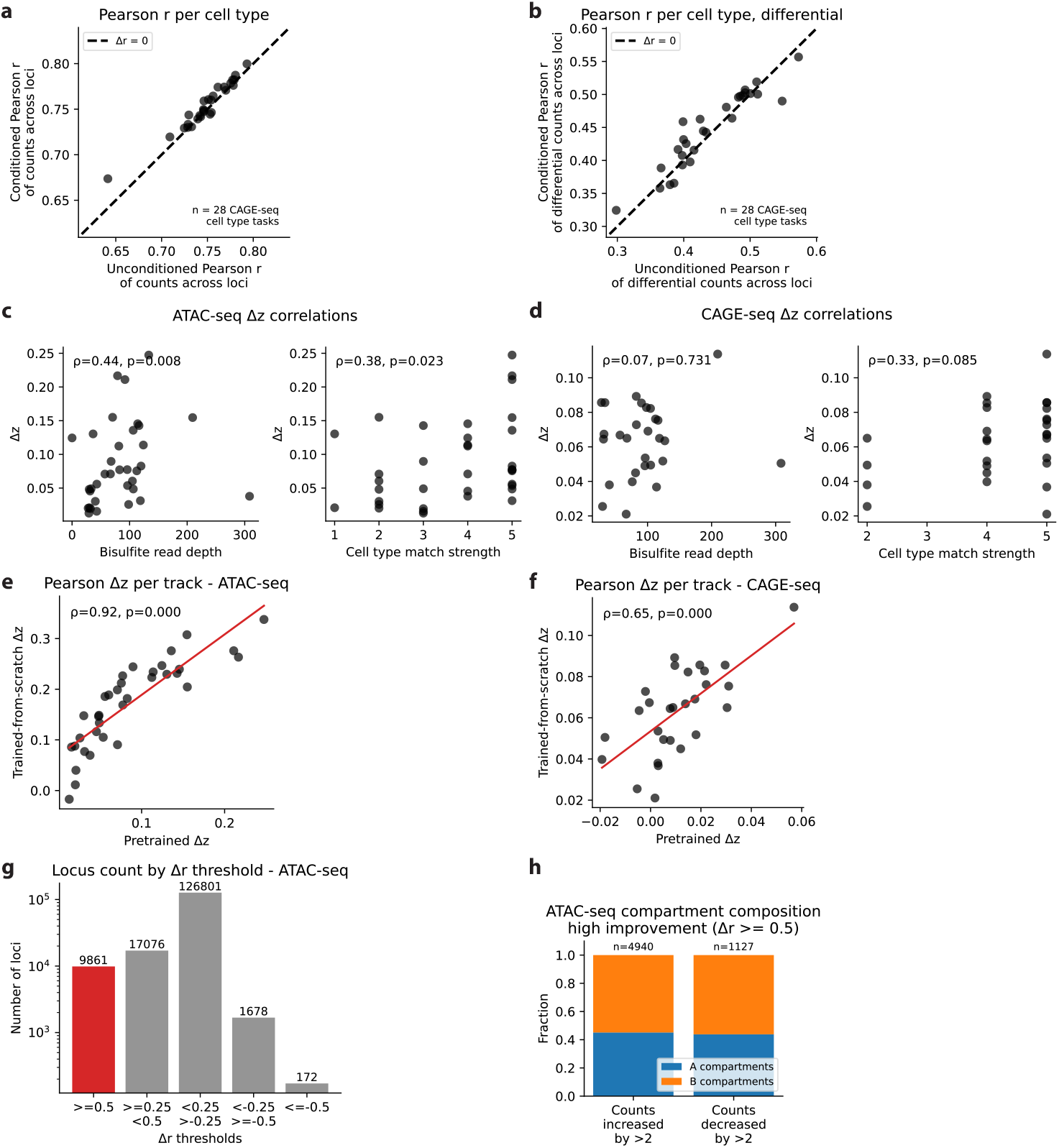
Cell type taskwise improvements. **a**, Scatterplot comparison of MethylSeqNet conditioned vs unconditioned prediction-measurement Pearson correlation for CAGE-seq test set by cell type. Each point represents one cell type/task, and the dashed line indicates equal performance. **b**, As in **a**, but for cell-type-differential Pearson correlation calculated from the difference from mean activity across cell types for the input sequence (Methods). **c**, ATAC-seq Fisher-z-transformed Pearson (Δz) conditioned-unconditioned by cell type correlated to WGBS read depth and to WGBS:ATAC-seq cell type match quality, showing Spearman correlation. **d**, As in **c**, but for CAGE-seq. **e**, ATAC-seq Fisher-z-transformed Pearson (Δz) conditioned-unconditioned by cell type for two MethylSeqNet configurations: a trained-from-scratch Basenji2 sequence encoder compared with the pretrained Borzoi encoder used for main-text analysis. **f**, As in **e**, but for CAGE-seq. **g**, Distribution of peak loci in terms of ATAC-seq per-locus cross-cell-type Pearson correlation change from Fig. 2c. **h**, Loci highlighted in **g** with per-locus cross-cell-type Pearson correlation increased by *>*0.5, assessed for all cell types with matched A and B compartment data from [31] and subset to cell types with *>*2 counts shift from unconditional to conditional predictions, as shown in Fig. 2d. Sites are localized into annotated A and B compartments for higher-activity-with-conditioning and lower-activity-with-conditioning peaks.

**Supplementary Fig. 4.**
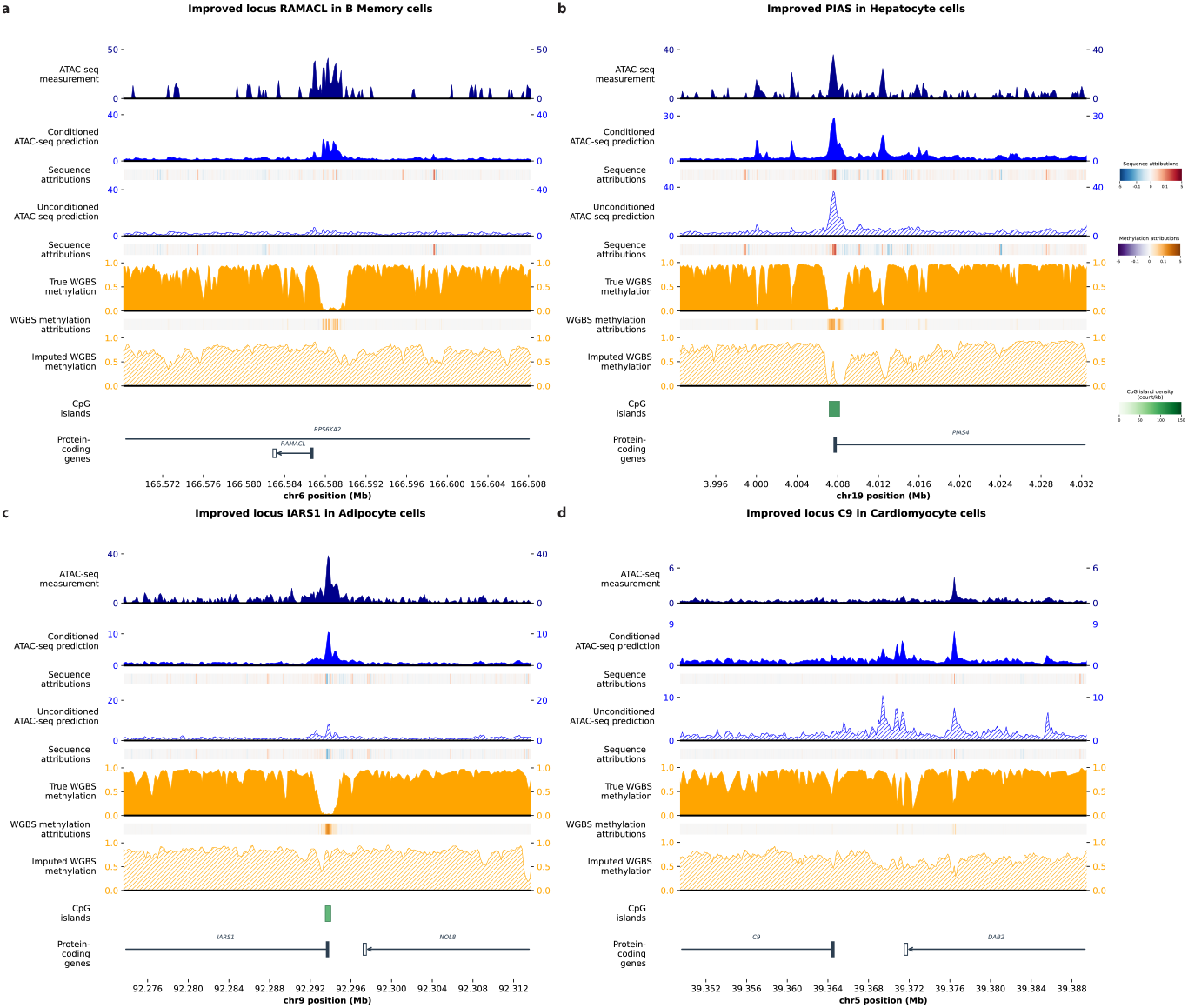
Visualizations of improved activity prediction at several loci. **a**,**b**,**c**,**d**, Examples of loci where MethylSeqNet improves accessibility prediction: *RAMACL* in B memory cells, *PIAS4* in Hepatocytes, *IARS1* in Adipocytes, and *C9* in Cardiomyocytes. We visualize genomic tracks for accessibility measurements, conditioned and unconditioned predictions from MethylSeqNet along with their respective sequence attributions calculated using integrated gradients, WGBS methylation measurements, MethylSeqNet imputed WGBS methylation, methylation attributions calculated using integrated gradients, and CpG island and gene annotations.

**Supplementary Fig. 5.**
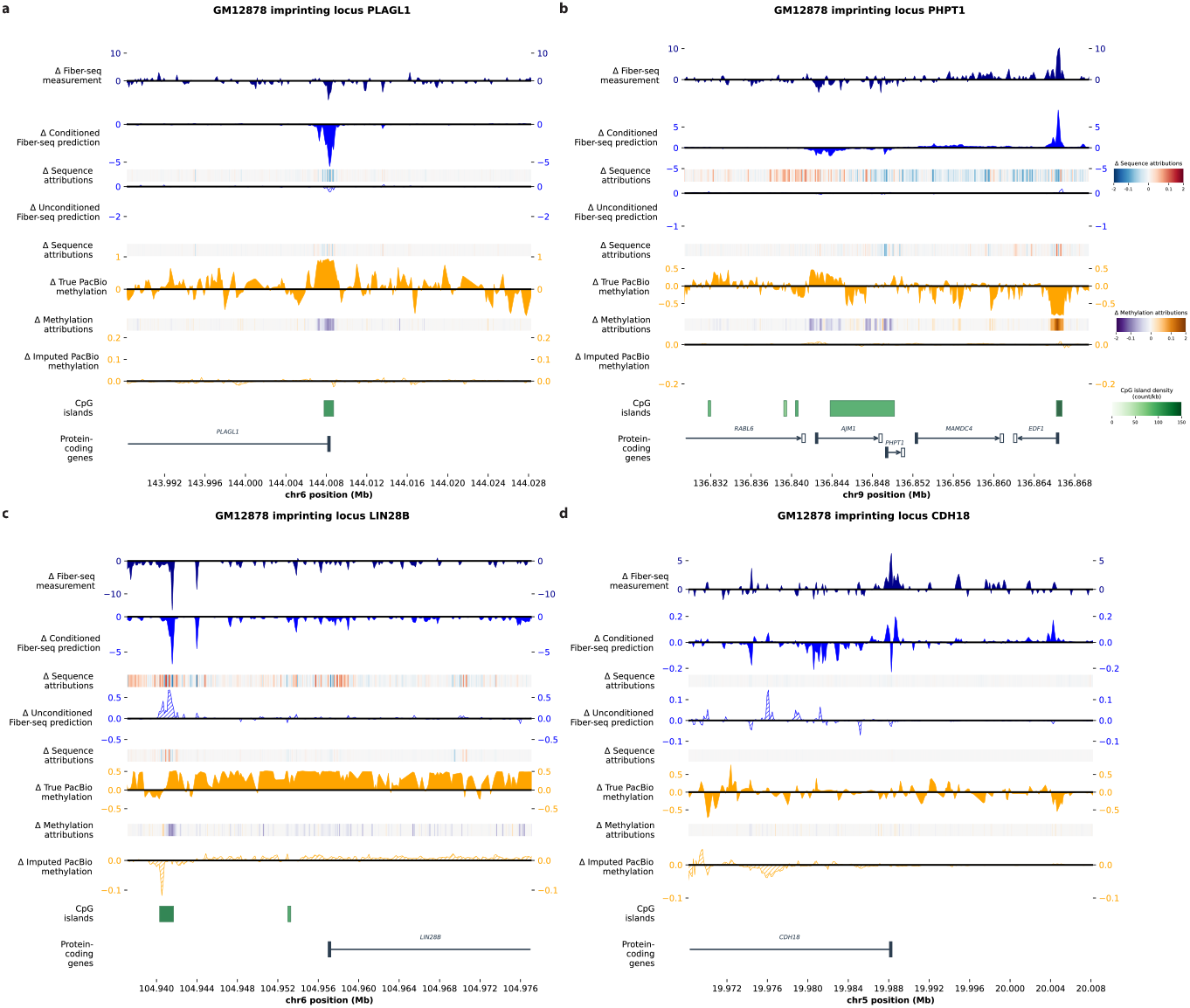
Visualizations of improved differential activity prediction at several imprinting loci. **a**,**b**,**c**,**d**, Examples of imprinting loci where MethylSeqNet improves haplotype-differential accessibility prediction: *PLAGL1, PHPT1, LIN28B*, and *CDH18*, all in GM12878 lymphoblastoid cells. We visualize haplotype-differential genomic tracks for accessibility measurements, conditioned and unconditioned predictions from MethylSeqNet along with their respective sequence attributions calculated using integrated gradients, PacBio methylation measurements, MethylSeqNet imputed PacBio methylation, methylation attributions calculated using integrated gradients, and CpG island and gene annotations.

**Supplementary Fig. 6.**
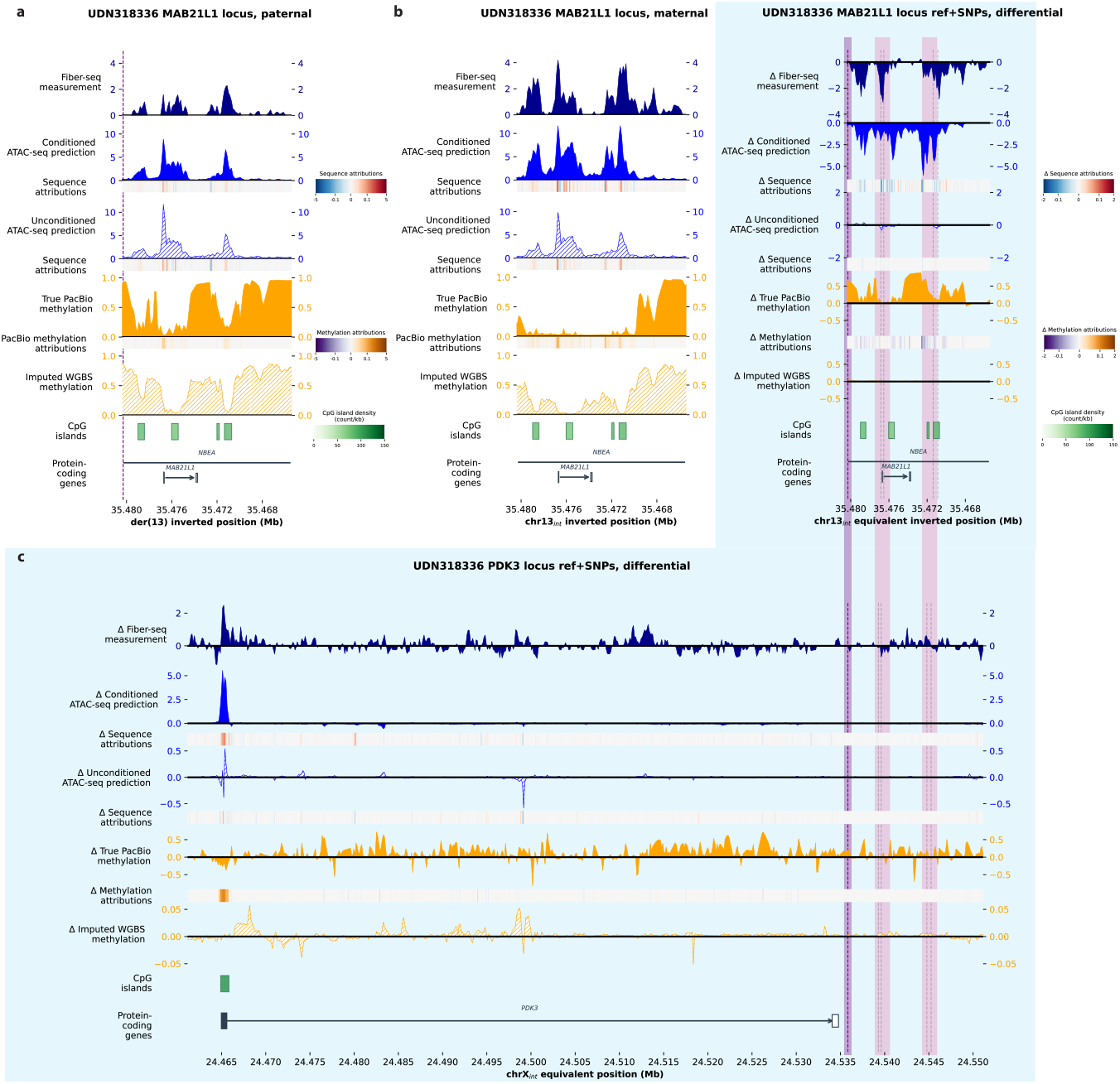
Undiagnosed disease patient loci. **a, b**, The *MAB21L1* locus shows large activity differences for paternal and maternal haplotypes. We visualize genomic tracks for accessibility measurements, conditioned and unconditioned activity predictions from MethylSeqNet along with their respective sequence attributions calculated using integrated gradients, PacBio methylation measurements, MethylSeqNet imputed methylation, methylation attributions calculated using integrated gradients, and CpG island and gene annotations. **c**, The *PDK3* and *MAB21L1* loci haplotype-differential activity predicted without inclusion of the large structural variant. Blue background indicates that the reference sequence with haplotype-specific SNPs was used, rather than the structural variant sequences. We visualize haplotype-differential genomic tracks for accessibility measurements, conditioned and unconditioned ATAC-seq predictions from MethylSeqNet’s neuron task along with their respective sequence attributions calculated using integrated gradients, PacBio methylation measurements, MethylSeqNet imputed WGBS methylation for neurons, methylation attributions calculated using integrated gradients, and CpG island and gene annotations. The fusion site (purple) and relevant promoters and enhancers (pink) are highlighted.

**Supplementary Fig. 7.**
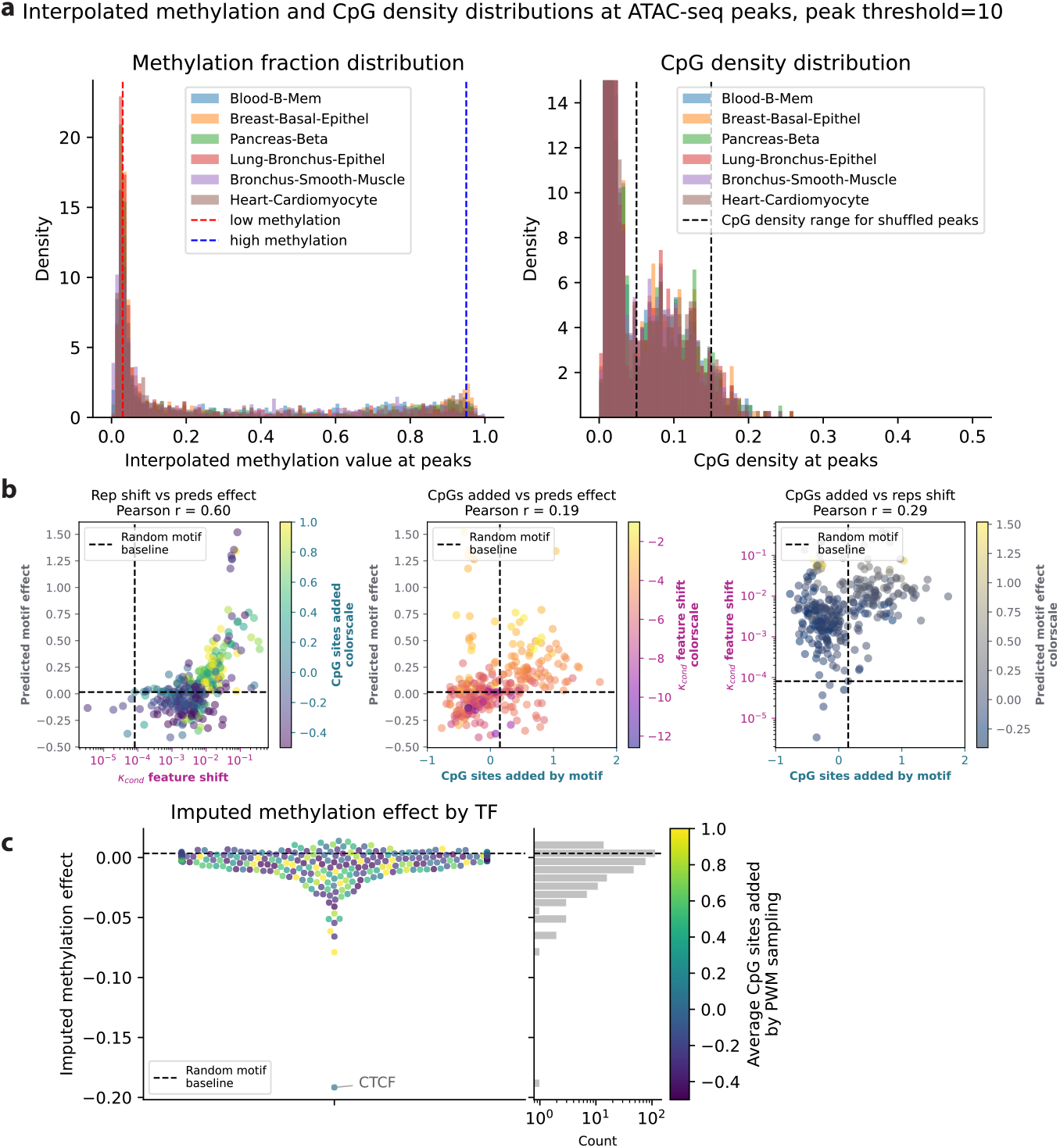
Further characterization of in silico motif insertion. **a**, Distributions of the accessibility peak methylation fraction distribution for WGBS when interpolated between CpG sites (left), and of the accessibility peak CpG density (right), for representative cell types. Dashed lines indicate values and ranges used for motif insertion. **b**, Scatterplots illustrate pairwise comparisons between three variables: chromatin accessibility TF motif effect, conditional feature shift, and average number of CpG sites added by a sampled position-weight matrix for that TF. The random motif baseline indicates values computed for a uniform random sampled motif. **c**, Swarmplot and corresponding histogram of the TF motif effect on imputed methylation predictions, showing CTCF as a strong outlier. Points are colored by the average number of CpG sites added by a sampled position-weight matrix for that TF.

**Supplementary Fig. 8.**
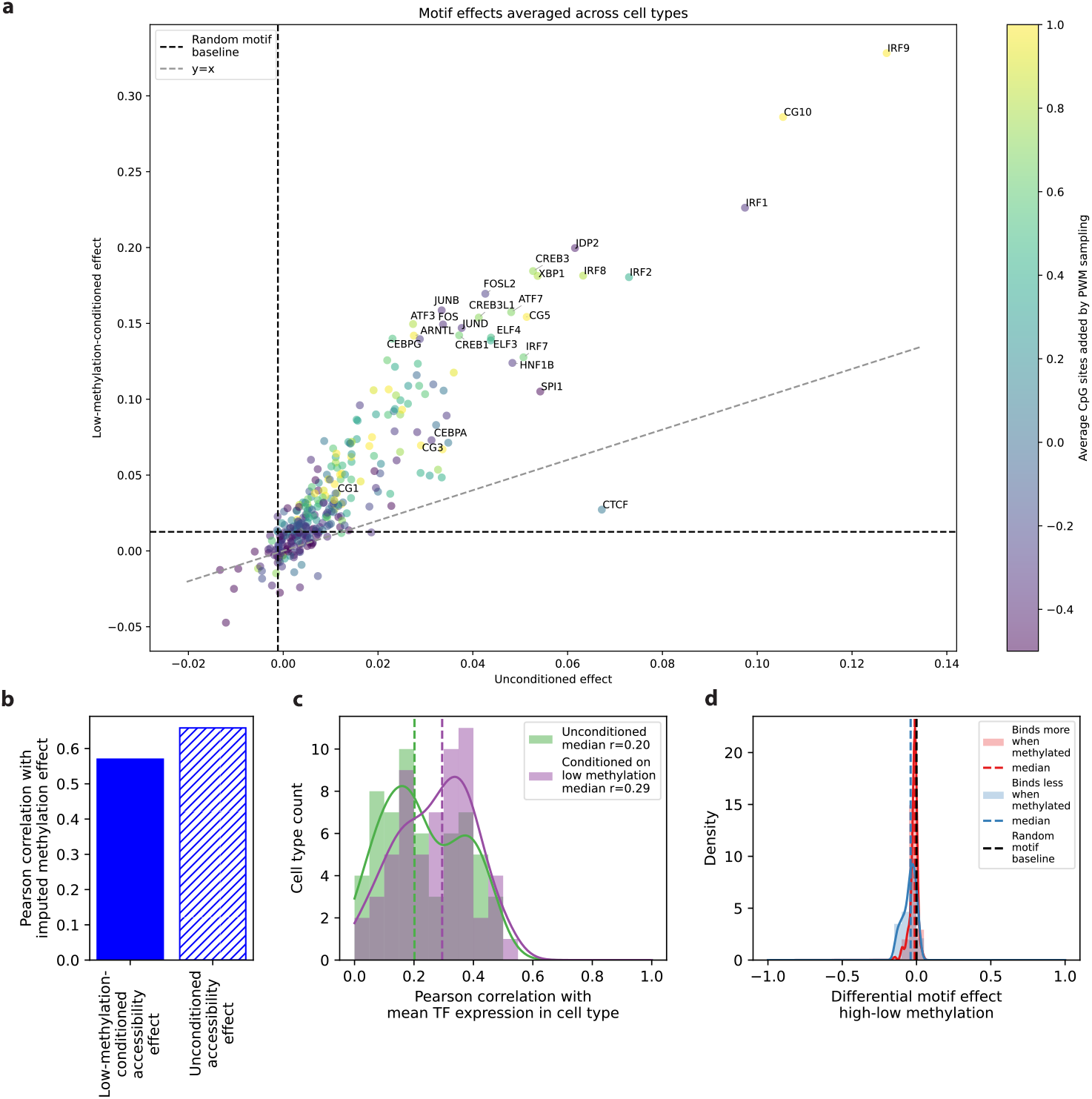
CAGE-seq TF motif effects from motif insertion analysis. **a**, Scatterplot of low-methylation-conditioned CAGE TF motif effect against unconditioned CAGE TF motif effect for all 301 TFs studied, averaged across all cell types. Points are colored by the average number of CpG sites added by a sampled position-weight matrix for that TF (the scale is truncated to exclude the all-CpG controls CG10, CG5, etc.), and several of the highest-effect TFs are highlighted. Black horizontal and vertical dashed lines indicate the calculated motif effect of a uniform random motif of length 10 for low methylation and unconditioned cases, and the gray dashed line indicates *y* = *x*. **b**, Correlation of the CAGE TF motif effects calculated using imputed methylation with those calculated using low methylation (left, solid) and no methylation conditioning (right, striped). **c**, Histograms of Pearson correlations per cell type between expression TF motif effect and corresponding TF expression in each cell type, plotted for low-methylation-conditioned vs unconditioned predictions. **d**, Histograms of differential CAGE TF motif effects (calculated as high methylation effect minus low methylation effect) for TFs known to bind more with methylation (red) vs those known to bind less (blue).

**Supplementary Fig. 9.**
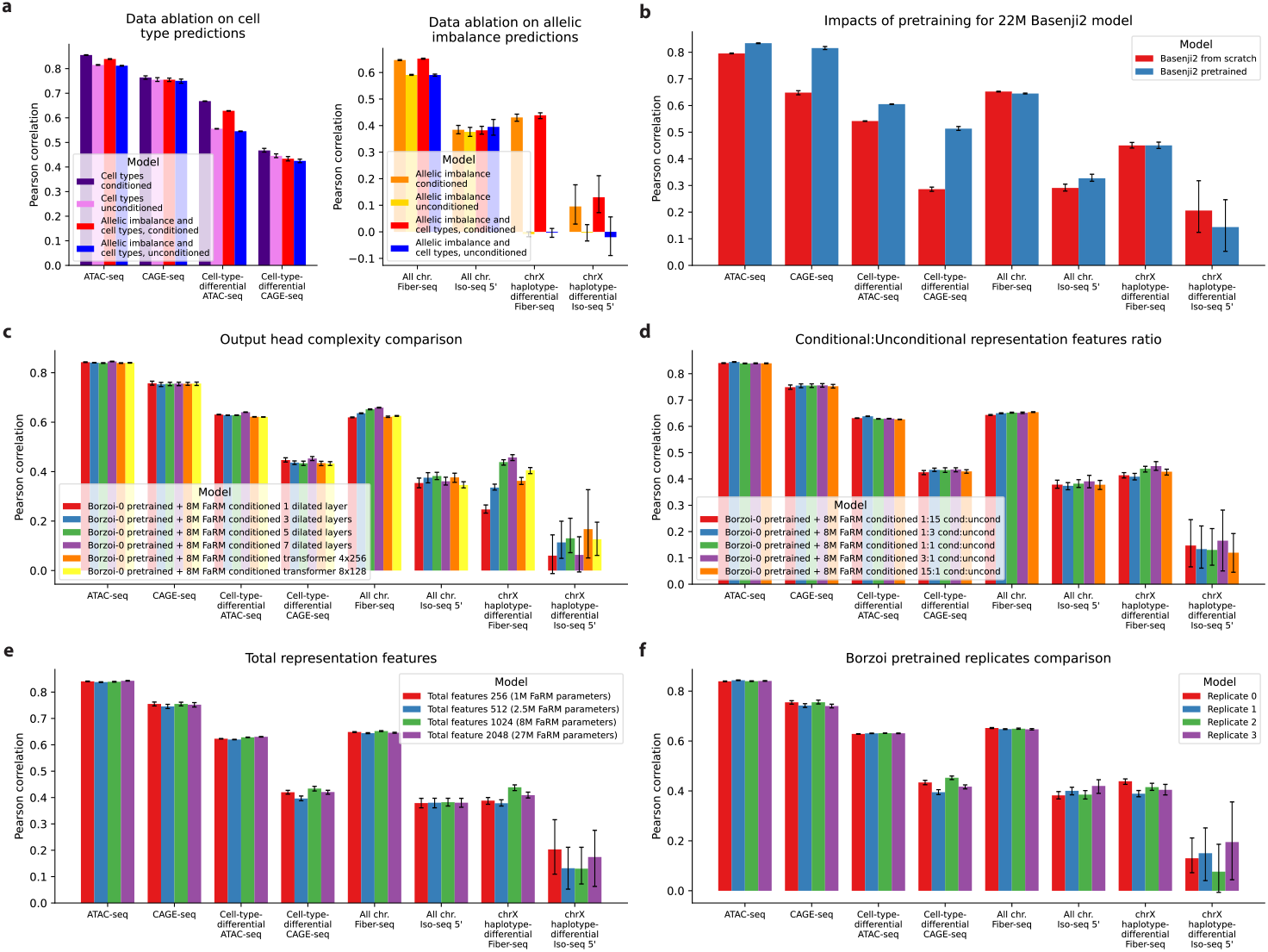
Performance comparisons of model ablations. **a**, Performance comparison from training conditioned and unconditioned MethylSeqNet on cell type and allelic imbalance data ablations, for the assay-relevant subsets of the data and output tasks. 95% confidence intervals from 100 bootstrap resamples (N = full test set subset size, with replacement) are shown. **b**, As in **a** but comparing Basenji2 trained from scratch vs pretrained in a MethylSeqNet architecture for various subsets of the data and output tasks, including both standard activity prediction and differential activity prediction between two sequences. **c**, As in **b** but comparing output (prediction) head architectures with increasing numbers of dilated convolution layers and two transformer layer configurations. **d**, As in **b** but comparing different ratios of conditional to unconditional features. **e**, As in **b** but comparing total representation features (conditional and unconditional). **f**, As in **b** but comparing the 4 pretrained Borzoi replicates.

